# Exploratory behavior is associated with microhabitat and evolutionary radiation in Lake Malawi cichlids

**DOI:** 10.1101/525378

**Authors:** Zachary V. Johnson, Emily C. Moore, Ryan Y. Wong, John R. Godwin, Jeffrey T. Streelman, Reade B. Roberts

## Abstract

Encountering and adaptively responding to unfamiliar or novel stimuli is a fundamental challenge facing animals and is linked to fitness. Behavioral responses to novel stimuli can differ strongly between closely related species; however, the ecological and evolutionary factors underlying these differences are not well understood, in part because most comparative investigations have focused on only two species. In this study, we investigate behavioral responses to novel environments, or exploratory behaviors, sampling from a total of 20 species in a previously untested vertebrate system, Lake Malawi cichlid fishes, which comprises hundreds of phenotypically diverse species that have diverged in the past one million years. We show generally conserved behavioral response patterns to different types of environmental stimuli in Lake Malawi cichlids, spanning multiple assays and paralleling other teleost and rodent lineages. Next, we demonstrate that more specific dimensions of exploratory behavior vary strongly among Lake Malawi cichlids, and that a large proportion of this variation is explained by species differences. We further show that species differences in open field behaviors are explained by microhabitat and by a major evolutionary split between the mbuna and benthic/utaka radiations in Lake Malawi. Lastly, we track some individuals across a subset of behavioral assays and show that patterns of behavioral covariation across contexts are characteristic of modular complex traits. Taken together, our results tie ecology and evolution to natural behavioral variation, and highlight Lake Malawi cichlids as a powerful system for understanding the biological basis of exploratory behaviors.

## 1. Introduction

Responding to unfamiliar or novel stimuli is a fundamental aspect of animal life that has important implications for fitness; how individuals behave during initial encounters with unfamiliar environments, predators, potential mating partners or competitors, and food resources can directly impact survival and reproduction (Dingemanse, Both, Drent et al., 2004; Ferrari, McCormick, Meekan et al., 2015; Lapiedra, Schoener, Leal et al., 2018; Smith & Blumstein, 2008). As a result, different behavioral responses to different types of novel stimuli may be favored in different ecological contexts.

Exposure to unfamiliar environments has been shown to trigger conserved physiological, neural, and behavioral responses in teleost fishes and mammals. In both zebrafish and mice, exposure to an unfamiliar open environment induces cortisol release (Gross, Santarelli, Brunner et al., 2000; Kysil, Meshalkina, Frick et al., 2017; R. Y. Wong, French, & Russ, 2019) and c-fos expression in the basolateral amygdala (in zebrafish the medial dorsal telencephalon, the putative teleost homologue) (Hale, Hay-Schmidt, Mikkelsen et al., 2008; Lau, Mathur, Gould et al., 2011; O’Connell & Hofmann, 2011). When most fish or rodent species are introduced to a large open field environment, they initially avoid the exposed center region and remain close to the outer edges and corners (thigmotaxis) (Champagne, Hoefnagels, de Kloet et al., 2010; Seibenhener & Wooten, 2015; Treit & Fundytus, 1988; Webster, Baumgardner, & Dewsbury, 1979; Wilson, Vacek, Lanier et al., 1976). Humans also have been shown to exhibit thigmotaxis in a natural “open field” environment, and patients with severe anxiety symptoms exhibit stronger thigmotaxis (Walz, Mühlberger, & Pauli, 2016).

Other types of novel stimuli elicit similar behavioral responses in teleosts and rodents. For example, when introduced to a “light-dark” chamber in which one half is brightly illuminated and the other half is dark, mice, rats, and many species of fish spend more time in the dark half (scototaxis) (Arrant, Schramm-Sapyta, & Kuhn, 2013; Dereje, Sawyer, Oxendine et al., 2012; C. Maximino, Marques de Brito, Dias et al., 2010; Takao & Miyakawa, 2006). In some cases, the same pharmacological compounds (e.g. fluoxetine and buspirone, both of which modulate serotonin signaling) decrease “avoidance” of the more open or exposed region of novel environments in both zebrafish and mice, and these compounds have also been used to treat psychiatric symptoms such as anxiety, panic, and/or depression in humans (Angelis, 1996; Dulawa, Holick, Gundersen et al., 2004; C. Maximino, A. W. da Silva, A. Gouveia, Jr. et al., 2011; C. Maximino, A. W. B. da Silva, A. Gouveia et al., 2011; Simon, Dupuis, & Costentin, 1994; R. Y. Wong, Oxendine, & Godwin, 2013).

Although broad patterns may be conserved, within taxa some species exhibit atypical and divergent behavioral responses to novel stimuli. For example, a comparative study of 12 muroid rodent species found while most species displayed the generally conserved pattern of thigmotaxis in the open field test, cotton mice (*Peromyscus gossypinus*) did not, making more entries into the center region than other parts of the arena (Wilson, Vacek, Lanier et al., 1976). Indeed, many dimensions of behavioral responses to novel stimuli can vary strongly between closely-related species (Reale, Reader, Sol et al., 2007); however, the factors underlying these examples of natural behavioral divergence are not well resolved.

Large scale comparative studies are a powerful strategy for identifying evolutionary and/or ecological factors underlying species differences in behaviors. For example, habitat is related to open field behaviors in muroid rodents, with semi-arboreal species exhibiting more exploratory phenotypes than species inhabiting desert and field habitats (Wilson, Vacek, Lanier et al., 1976). Similarly, a comparative study across 61 species of parrots showed that microhabitat explained species variation in responses to novel objects: species inhabiting intermediate habitats between the forest and the savannah more readily approached novel objects compared to species inhabiting more uniform savannah habitats (Greenberg, 2003; Greenberg & Mettke-hofmann, 2001; Claudia Mettke-Hofmann, Winkler, & Leisler, 2002). These and other data support the idea that ecological divergence may explain natural variation in exploratory behaviors among species. However, it is unclear if this model predicts behavioral differences in other species groups and in different vertebrate lineages, in part because many comparative studies have compared just two species (Garland & Adolph, 1994; Réale, Reader, Sol et al., 2007). Furthermore, different behavioral assays and testing parameters have been used across many studies, making it difficult to identify common factors that explain specific dimensions of behavior. To better elucidate relationships between ecological factors, such as microhabitat, and species differences in exploratory behaviors, large comparative studies in new vertebrate systems are needed.

Lake Malawi cichlid fishes are well-suited for comparative investigations of phenotypic variation, and have attracted the attention of evolutionary biologists for more than a century (R. C. Albertson, Markert, Danley et al., 1999; Johnson & Young, 2018; Rupp & Hulsey, 2014; Ryan A. York & Fernald, 2017). These fishes have recently (within the past one million years) undergone explosive speciation, diversifying through multiple major evolutionary radiations into an estimated 500-1000 species that vary in morphology, coloration, diet, habitat preference, and behavior (Brawand, Wagner, Li et al., 2014; Kocher, 2004; Malinsky, Svardal, Tyers et al., 2018). Within Lake Malawi, ecological conditions vary across small spatial scales, resulting in diverse species occupying different microhabitats while living in close geographic proximity. For example, although many species can be grouped into two canonical ecotypes, rock-dwelling and sand-dwelling (Kocher, 2004), a large number of species occupy the intermediate habitat, or the interface between rocky and sandy substrate. Thus, the Lake Malawi species assemblage is an excellent system for studying relationships between evolution, ecology, and phenotypic variation.

Comparative studies in Lake Malawi cichlids have already generated insights into the evolution of complex traits. Ecological factors have been linked to species differences in jaw morphology and behaviors such as aggression and bower-building (R. Craig Albertson, 2008; Danley, 2011; Ryan A. York, Patil, Hulsey et al., 2015). Species differences in morphology, color patterning, sex determination, and bower building behavior have also been mapped to specific genomic loci (R. Craig Albertson, Streelman, & Kocher, 2003; Bloomquist, Parnell, Phillips et al., 2015; Conith, Hu, Conith et al., 2018; Kratochwil, Liang, Gerwin et al., 2018; Roberts, Ser, & Kocher, 2009; Ser, Roberts, & Kocher, 2010; R. A. York, Patil, Abdilleh et al., 2018).

Malawi cichlids are also an excellent system for studying phenotypic integration and modularity. Several studies have found modular patterns of phenotypic variation for complex traits, in particular traits that are thought to have played a central role in cichlid diversification, such as oral jaw morphology and color patterning (R. Craig Albertson, Powder, Hu et al., 2014; Parsons, Cooper, & Albertson, 2011). Briefly, evolutionary modularity and integration refer to distinct patterns of covariation among sets of traits across taxa, and they are thought to be related to trait evolvability (Raff & Raff, 2000; Wagner, Pavlicev, & Cheverud, 2007). For example, if the dimensions of different oral jaw bones are correlated in the same way across species, then they are considered to be evolutionarily integrated. In contrast, if they are uncorrelated or are correlated non-uniformly across taxa, they are more modular and are generally considered to be more evolvable; however, integration does not necessarily suggest a constraint on evolvability, and patterns of covariation by themselves are not sufficient for proving lesser or greater evolutionary potential (Armbruster, Pélabon, Bolstad et al., 2014).

Although the Lake Malawi cichlid assemblage is an excellent system for large scale comparative investigations, few comparative behavioral studies have been conducted in this system. We aim to address this gap by investigating behavioral responses to novel environments using three classic assays: the novel tank test, the light-dark test, and the open field test (Stewart, Cachat, Wong et al., 2011; Stewart, Gaikwad, Kyzar et al., 2012). In each assay, we sample different subsets of species, ultimately spanning a total of 20 species and three Lake Malawi microhabitats: rock, sand, and rock/sand intermediate. We test the following hypotheses: (i) Malawi cichlids exhibit general responses to novel stimuli that are similar to other teleosts and other vertebrates; (ii) natural evolution has resulted in a high degree of phenotypic variance in exploratory behaviors among Lake Malawi cichlids; (iii) variation in exploratory behaviors is explained by divergence between species; (iv) species differences in exploratory behaviors are associated with differences in microhabitat and with major evolutionary radiations in Malawi cichlids; and (v) like other complex traits in this species assemblage, exploratory behaviors are modular.

## 2. Methods

### 2.1 Subjects

Subjects were maintained at two institutions, Georgia Institute of Technology (INSTITUTION 1) in Atlanta, GA and North Carolina State University (INSTITUTION 2) in Raleigh, NC. Both institutions house laboratory cichlid lines derived from wild-caught animals collected in Lake Malawi and subsequently cultured in laboratory aquarium environments. All subjects were fertilized and raised in laboratory aquariums. Similar housing and husbandry conditions were maintained at both institutions: (i) age- and size-matched individuals were socially housed in mixed-sex tanks at similar densities (ranging between 0.67-1.33 cm of fish/liter) and co-cultured as necessary to reduce aggression; (ii) all tanks were maintained on a central recirculating system, and thus water conditions across tanks were the same; (iii) water temperatures at both institutions ranged between 27-28°C, pH between 7.8-8.2, and conductivity between 230-260 uS; (iv) room temperatures at both institutions ranged from 26.5-28.0°C and humidity was maintained at approximately 40%; (v) all species were housed in glass-bottom aquariums with some structural enrichment (air stones, plastic/PVC pipes, ceramic tiles, and/or terracotta pots); and (vi) 12:12 hour light:dark cycles were maintained with transitional dim light periods.

INSTITUTION 1 animals were maintained in the Engineered Biosystems Building cichlid aquaculture facilities at INSTITUTION 1 in accordance with the Institutional Animal Care and Use Committee (IACUC) guidelines (protocol numbers A100028 and A100029). Subjects were housed on a 12:12-hour light:dark cycle with full lights on between 8am-6pm Eastern Standard Time (EST) and dim lights on for 60 minutes between the light-dark transition (7am-8am and 6pm-7pm EST). All subjects were housed in 190-liter or 95-liter glass tanks measuring 92 cm (long) x 46 cm (wide) x 42 cm (high) or 46 cm (long) x 46 cm (wide) x 42 cm (high), respectively, and fed daily (Spirulina Flake; Aquatic Ecosystems). Male and female subadults (age 90-180 days) were analyzed in the novel tank test and light-dark test (described below), and male and female reproductive adults (>180 days) were tested in the open field test (described below).

INSTITUTION 2 animals were maintained in the INSTITUTION 2 Roberts Lab cichlid aquaculture facility in Raleigh, NC. Subjects were housed on a 12:12-hour light:dark cycle with dim lights on for 15 minutes during the light-dark transition periods, and were fed daily (Best Flake 70% Vegetable/30% Brine mix; Worldwide Aquatics). All experiments were conducted under the approval of the Institutional Animal Care and Use Committee (IACUC) guidelines (protocol number 14-138-O). For all thirteen INSTITUTION 2 species tested in the open field test, subjects were housed in 189-liter or 473-liter tanks measuring 92 cm (long) x 47 cm (wide) x 48 cm (tall) or 184 cm (long) x 47 cm (wide) x 60 cm (tall), respectively, and were tested as male and female reproductive adults (>180 days).

### 2.2 Animal welfare

At both institutions, the utmost care was taken to minimize stress from handling and housing, both in general husbandry and during behavioral experiments. Outside of behavioral testing, fish were communally housed to provide social interaction, and monitored carefully to ensure that animals were not the target of extreme aggression. Cichlids naturally establish social hierarchies in the home tank (and in the wild) through aggression. In cases of animals being repeatedly targeted with aggression in the home tank, individuals were transferred to a different social tank to maximize the safety of all animals throughout the course of the study. During behavioral experiments, fish were gently netted out of their home tanks by an experienced handler and carefully moved to reduce stress as much as possible. Transfer containers were covered by nets to reduce stress, as well.

### 2.3 Behavioral assays

A total of 455 subjects spanning 20 Lake Malawi cichlid species were tested in one or more assays that are well-established and designed to measure exploratory behaviors in teleosts. Pilot data indicated strong effects of species but no effects of sex on exploratory behaviors across multiple assays. Based on these data, subjects for the present study were sampled randomly from mixed sex tanks but were not euthanized and dissected to determine gonadal sex, with the exception that visually identified dominant males were sampled at a proportion consistent with the composition of the home tank, and maternal mouthbrooding females were not sampled. All assays were performed between 10:00 and 16:00 Eastern Standard Time EST. Each assay is described in detail by institution (INSTITUTION 1 and INSTITUTION 2), species, sample size, and experimental design in the following sections.

#### Assays by test site

The novel tank and light-dark tests were conducted at INSTITUTION 1 only. 110 subjects from eight species were tested in the novel tank test; 67 of these subjects were also tested in the light-dark test, and four additional subjects were tested exclusively in the light-dark test (see Supplementary Tables 1, 2, and 5 for sample sizes by species). The open field test was then conducted across a larger species and subject pool spanning both INSTITUTION 1 and INSTITUTION 2. For the open field test, 341 subjects from 19 species were tested: 227 subjects from 13 species at INSTITUTION 2, and 113 subjects from seven species at INSTITUTION 1, with one species (*Labeotropheus fuelleborni)* tested at both institutions (See Supplementary Table 3 for sample sizes by species).

To assess phenotypic integration versus modularity of exploratory behaviors, correlated behaviors across novel contexts were measured with Modularity Modular Clustering analysis (MMC; described below), a well-established technique that was developed for analyzing patterns of covariation in gene expression data (Stone & Ayroles, 2009). Because measurement of behavioral covariance is a novel application of MMC, we first confirmed that it could reveal clusters of behavioral covariation across assays. To do this, we used MMC to analyze a previously published control zebrafish dataset, in which 99 subjects from three selection lines were tested across a battery of behavioral assays and correlated behaviors across contexts were found (Ryan Y Wong, Perrin, Oxendine et al., 2012). The second dataset included 67 subjects from eight Malawi cichlid species that were tracked across the novel tank test and light-dark test at INSTITUTION 1 (Supplementary Table 4). The third dataset included 70 subjects from five Malawi cichlid species that were tracked across a rectangular open field test, resident-intruder test, and novel object test as part of a separate set of experiments at INSTITUTION 2 (see “Supplementary Methods and Results” and Supplementary Table 4).

#### Novel tank test

The novel tank test is a classic assay designed to measure exploration of a tall and narrow transparent tank, with primary focus on exploration of the upper half (Fig. 1A-B) (Cachat, Stewart, Grossman et al., 2010; Egan, Bergner, Hart et al., 2009; K. Wong, Elegante, Bartels et al., 2010). Individual subadult subjects (90-180 days; 1.75-2.5 cm standard length, SL) spanning eight species were collected between 11:00-15:00 Eastern Standard Time from their home tank, transferred to a 300 mL holding beaker, and habituated for 30 minutes prior to behavioral testing. Water for both habituation beakers and test tanks was collected from a recirculating aquaculture system supplying all home tanks, ensuring that water was consistent across the home tank, transfer, habituation, and testing environments. Following habituation, subjects were introduced to a plastic 1.8-L novel tank (Aquaneering; 29.7 cm long x 7.5 cm wide 15.2 cm high) and were side-view video recorded for 6 minutes using a GoPro Hero4 camera. Species composition was counterbalanced across trials to control for potential effects of testing round. EthoVision (Noldus) software was used to analyze time spent in the top half, entries/exits to and from the top half, latency to enter top half, average distance from the bottom and corners, and total distance traveled.

**Figure 1.**
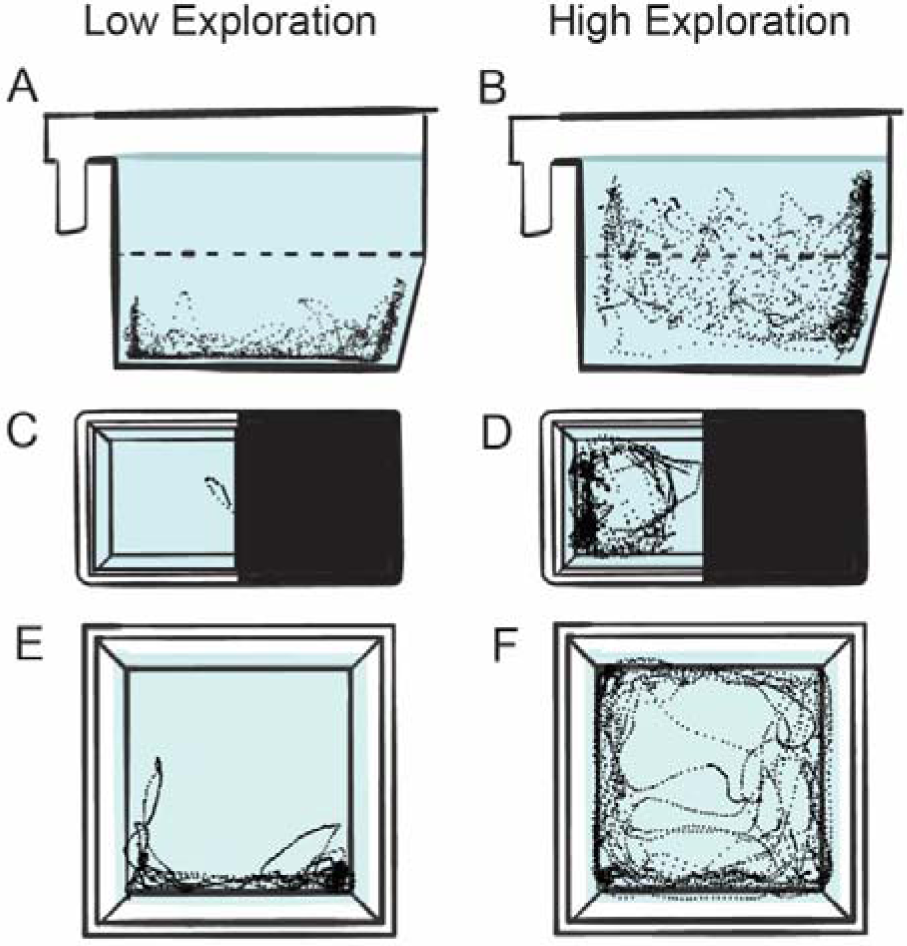
Lake Malawi cichlids exhibit high phenotypic variation in exploratory behaviors. Behavioral variation illustrated by representative traces from the four behavioral assays used in this study, the novel tank test (A-B), light-dark test (C-D), and open field test (E-F). Individual points illustrate the position of the subject in the arena at a single timepoint. A high degree of phenotypic variance was observed across assays, ranging from stereotypically low exploratory phenotypes (A,C,E) to high exploratory phenotypes (B,D,F). For each assay, the schematic reflects the camera angle from which video recordings were collected for each trial.

#### Light-dark test

In the light-dark test, subjects can freely move between an opaque black chamber and a backlit semi-opaque white chamber (Fig. 1C-D). As in rodents, this assay is designed to investigate place preferences between a dark versus illuminated environment, and exploration of an unfamiliar illuminated environment (Bourin & Hascoet, 2003; Champagne, Hoefnagels, de Kloet et al., 2010; Hascoet, Bourin, & Nic Dhonnchadha, 2001). Individual subadult subjects (90-180 days; 1.75-2.5 cm SL) from all eight tested species were transferred to a 300 mL beaker of water and habituated for 30 minutes prior to testing. All water was collected from the same recirculating aquaculture system (described above). Following habituation, subjects were first introduced to a 6.5 cm x 7.5 cm habituation chamber (half white, half black) within the larger custom built acrylic light-dark tank (half white, half black; 24 cm long x 6.5 cm wide x 16.5 cm high). Individual subjects habituated for 5 minutes in the central habituation chamber, at which point two inserts were simultaneously removed, allowing subjects to swim freely throughout the entirety of the light-dark tank. Species composition was counterbalanced across trials to control for potential effects of testing round. All subjects were top-down video recorded for 6 minutes using a GoPro Hero4 camera. EthoVision (Noldus) software was used to analyze time spent in the light versus dark halves, as well as latency to enter, number of entries, total time spent, and total distance traveled in the light half.

#### Open field test

The open field test for teleosts is similar in design to the open field test used in mice and other rodents, in which subjects are allowed to move freely throughout a large open arena (Champagne, Hoefnagels, de Kloet et al., 2010; Seibenhener & Wooten, 2015). For teleosts, vertical motion is restricted by shallow water depth, and the test is thus designed to measure behavioral responses to a large and open shallow water environment (Fig. 1E-F). For the present study, 19 species were analyzed in the open field test at two test sites (INSTITUTION 1 and INSTITUTION 2). MMC (described below) also included re-analysis of a separate open field (and two additional behavioral assays) dataset collected as part of a different study (see (Emily C. Moore & Roberts, 2019)) from five species under different parameters (described below) at INSTITUTION 2 (see Supplementary Methods and Results and Supplementary Table 4).

All subjects were gently netted from their home tank and placed in the center of a white, opaque container filled with aquaculture system water at shallow depths to restrict vertical movement. At both institutions, larger subjects exceeding 4.5 cm standard length (SL) were introduced to a 49.6 cm-wide square arena filled to a depth of 15 cm, while smaller subjects ranging from 2.5-4.5 cm SL were introduced to a 25.5 cm-wide square arena filled to a depth of 10 cm.

For all open field trials, tank water was replaced between every subject. Video recordings were taken for 5.5 minutes from an overhead position. The first 10 seconds of the video files were trimmed (Quicktime Player 7) to remove footage of subject placement. Videos were processed at 10 frames per second (fps) using C-trax 0.5.4 (Branson et al. 2009) to generate X, Y coordinates of fish position in arena. Custom scripts were used to generate position and speed in the arena (R v3.3.1). For place analysis, the arena was divided into a grid of 16 squares, with the outer ring of squares forming the “peripheral” regions, the central four squares forming the “center” region, and the four corner squares forming the “corner” regions.

### 2.4 Designations of microhabitat, evolutionary radiation, and genus

Previous genomic analyses suggest that Lake Malawi cichlids have diversified through multiple major evolutionary radiations of (i) pelagic species, (ii) shallow/deep benthic and “utaka” species, and (iii) “mbuna” species (Malinsky, Svardal, Tyers et al., 2018). The species sampled in the present study represented the latter two radiations (shallow/deep benthic and utaka, B/U; and mbuna). These radiations are well-characterized, and designations for evolutionary radiation as well as genus were made according to Konings (Konings, 2007). Microhabitat designations (rock, sand, or intermediate) for each species were made according to Ribbink et al. and Konings (Konings, 2007; Ribbink, Marsh, Marsh et al., 1983).

### 2.5 Statistics

All statistical analyses were performed in R (R v3.3.1 or later) unless otherwise specified.

#### Place biases across assays

To measure general place biases between zones in the novel tank and light-dark tests across species, a linear regression model with time spent as the outcome variable, and zone (e.g. top vs. bottom) and species as categorical predictor variables, was fit to the data.

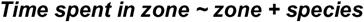

Because the open field test was performed at two test sites using two arena sizes, these factors were added to the model as categorical variables, and time spent in central versus peripheral regions were analyzed:

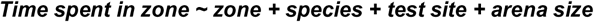

Within each species, paired t-tests were used to test the significance of differences in time spent in different zones.

#### Phenotypic variance in exploratory behaviors

Phenotypic variance was calculated for novel tank behaviors in both Malawi cichlids and high- and low-exploratory zebrafish strains. These included total time spent in the top half, number of entries into the top half, and latency to enter the top half. For each behavior, the overall variance observed among Lake Malawi cichlids was calculated, and the overall variance observed among high- and low-exploratory zebrafish strains was calculated. An F-test was used to quantify the difference between behavioral variance among Malawi cichlids and behavioral variance among high- and low-exploratory zebrafish strains.

#### Species differences in exploratory behavior

One-way ANOVA was used to test for species differences in behaviors. Effect size (Eta-squared) was calculated by dividing the individual effects’ sum of squares by the total sum of squares.

#### Associations among microhabitat, radiation, and open field behaviors

Associations between microhabitat and behavior were assessed through linear mixed effects models using the “lme4” package in R. Each behavior of interest was designated as the outcome variable, microhabitat and evolutionary radiation (mbuna vs. B/U) as fixed effects, species nested within genus as a random effect, and both arena size and lab as random effects. In this model microhabitat and evolutionary radiation directly competed to explain variance in exploratory behavior, controlling for variance explained by other phylogenetic factors and batch-like effects such as arena size and test site. This model was used to test six open field behavioral metrics, including time spent in the corners, entries into the corners, time spent in the center, entries into the center, total distance traveled, and change in speed over time. The model was organized as follows, (with bold italicized terms representing fixed effects, and non-bold italicized terms representing random effects, with nested terms in parentheses):

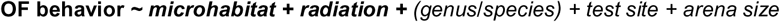

Because the mbuna radiation tends to inhabit rock microhabitats, and the B/U radiation tends to inhabit sand microhabitats, the fixed effects (radiation and microhabitat) in the above model were correlated, potentially masking additional relationships between microhabitat, evolutionary radiation, and exploratory behaviors. To further disentangle the relationships between the intermediate microhabitat, evolutionary radiation, and exploratory behavior, we applied a second model in which the original microhabitat term (rock, sand, or intermediate) was simplified into an intermediate (versus non-intermediate) term. This model thus allowed us to test how divergence into the intermediate microhabitat was associated with exploratory behaviors, controlling for variation explained by evolutionary radiation. To account for the possibility that divergence into the intermediate habitat may be differentially related to exploratory behaviors in the mbuna versus B/U radiations, we also included an intermediate*radiation interaction term:

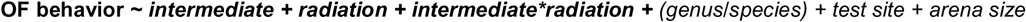

#### Associations among microhabitat, radiation, and novel tank corner behavior

To test whether mbuna rock-dwellers and B/U sand-dwellers exhibited differences in novel tank behavior, a simpler model was used (all species came from a unique genus, and all subjects were tested in identical tanks at the same test site). Notably, because all INSTITUTION 1 mbuna species inhabit rock habitats, and all INSTITUTION 1 B/U species inhabit sand habitats, “radiation” and “microhabitat” could be interchanged in the model with identical results:

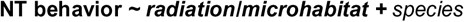

For all linear mixed effects models, estimates for fixed effects were calculated by maximum likelihood estimation using the ‘lme4’ package in R, and significance for fixed effects was calculated using Satterthwaite approximation through the ‘lmerTest’ package and the anova function in R. Estimates of pairwise differences between levels for each fixed effect were calculated using estimated marginal means (least squared means), and the significance of these differences were determined using Satterthwaite approximation corrected for multiple comparison families with Tukey’s adjustment, using the ‘emmeans’ and ‘multcomp’ packages in R.

To analyze movement in the open field test over time, the numbers of slow or stopped instances were summed over each minute, and 60-second bins were used as the input for a repeated measures MANOVA. Time points within individuals were analyzed at one level, differences between microhabitat were analyzed at an additional level, microhabitat*time was included as an interaction term, and additional terms were included to control for test site and arena size. The overall change in velocity (average velocity in minute 1 – average velocity in minute 5) throughout the assay was analyzed with an ANOVA by microhabitat (a positive value indicates that the subject swam faster at the start of the assay, and a negative value indicates the subject swam faster at the end of the assay).

#### Behavioral modularity

To analyze behavioral correlations within and across assays, we performed Modulated Modularity Clustering (MMC) analysis (Stone & Ayroles, 2009). This test identifies clusters of covariance in multivariate data. Although this method was developed to analyze gene expression data, it is effective for any large, multivariate datasets where many phenotypes have been measured across a large sample of subjects. To demonstrate as a proof-of-principle that MMC analysis can reveal behavioral correlations across assays, we re-analyzed a previously published zebrafish dataset in which individuals from selectively bred high- and low-exploratory strains were tracked across multiple assays and behavioral correlations across assays were found (Ryan Y Wong, Perrin, Oxendine et al., 2012). We then separately performed MMC on two independent Lake Malawi cichlid datasets: an INSTITUTION 1 dataset in which individuals were tracked across the novel tank and light-dark tests (Supplementary Table 4), and an INSTITUTION 2 dataset to analyze behavioral modules across the open field, resident-intruder, and novel object tests (Supplementary Methods and Results; Supplementary Table 4). In all MMC analyses, each individual behavioral metric within each assay (such as speed, position, time spent in a specific zone, etc.) was included in the analysis. Since these assays are of different measurement types, Spearman rank-order correlation was used in place of Pearson’s correlation.

## 3. Results

### 3.1 Malawi cichlids exhibit consistent place biases across assays

The three novel environment assays used in this study have been used widely in teleosts, particularly in zebrafish, and variations of these tests are well-established in rodents (Champagne, Hoefnagels, de Kloet et al., 2010; Seibenhener & Wooten, 2015; Treit & Fundytus, 1988; Wilson, Vacek, Lanier et al., 1976). We first investigated how Lake Malawi cichlids respond to these novel environments by measuring their place biases between different zones (e.g. light half versus dark half). In general, Lake Malawi cichlids exhibited strong place biases for specific zones in all three novel environment assays, spending more time in the bottom half of the novel tank test, the dark half of the light-dark test, and the periphery of the open field test. The directions of the place biases were the same in all species tested, and were consistent with those expressed by other teleosts and rodents. More detailed results are organized by assay below:

#### Malawi cichlids spend more time in the bottom region in the novel tank test

Linear regression controlling for species revealed that Malawi cichlids generally expressed a strong place preference for the bottom half in the novel tank test (n=110; t=20.982; p<0.0001), spending an average of 307.5±6.1 seconds in the bottom half compared to 52.5±6.1 seconds in the top half. The direction of the preference was consistent across all species tested, and two-tailed paired t-tests showed that this preference was significant within each species (p<0.05 for all species tested, Supplementary Table 1). Notably, *post-hoc* Tukey’s HSD tests showed significant differences in the strength of the bias between *Mchenga conophoros*, a B/U sand-dwelling species, and all other species tested, with *Mchenga conophoros* spending significantly more time in the top half (Supplementary Figure 1A). More detailed results by species are shown in Supplementary Table 1.

#### Malawi cichlids spend more time in the dark region in the light-dark test

Malawi cichlids exhibited a strong place bias in the light-dark test (n=77; t=16.07; p<0.0001), spending more time in the dark half (an average of 283.2±8.9 seconds in the dark half versus 76.8±8.9 seconds in the light half). Detailed results are presented by species in Supplementary Table 2. Notably, one B/U sand-dwelling species, *Copadichromis virginalis*, did not exhibit a significant place bias between the light and dark zones (n=12; two-tailed paired t-test, p=0.46; Supplementary Table 2), and this differed significantly from several other species (Supplementary Figure 1B). Additional results are presented by species in Supplementary Table 2.

#### Malawi cichlids spend more time in peripheral regions in the open field test

Malawi cichlids spent more time in the peripheral regions of the open field test compared to the center region. Linear regression controlling for species, test site, and arena size showed a strong place bias between the central versus peripheral regions (n=340; t=89.24; p<0.0001); spending an average of 298.9±2.2 seconds in the periphery compared to 21.1±2.2 seconds in the center. Two-tailed paired t-tests revealed these differences to be significant in every species tested (p<0.05 for all species tested, Supplementary Table 3). Additional results are presented by species in Supplementary Table 3. Notably, the B/U intermediate species *Aulonocara baenschi* and the mbuna rock-dweller *Metriaclima mbenjii* spent significantly less time in corner regions compared to multiple other species (Supplementary Figure 1C).

### 3.2 Malawi cichlids exhibit high phenotypic variance in exploratory behaviors

We next investigated phenotypic variance in exploratory behaviors. For a frame of reference, we compared phenotypic variance among Lake Malawi cichlids and among three strains of zebrafish: two wild-derived strains that have been selectively bred for divergent exploratory behaviors and a common domesticated wild-type strain (AB). This analysis was restricted to novel tank behaviors, because the test parameters used in the present study were the same as those used in the zebrafish study. For time spent in the top half, Malawi cichlids collectively exhibited greater phenotypic variance compared to the high- and low-exploratory zebrafish strains (n=110 Malawi cichlid individuals from eight species, n=99 zebrafish from three selection lines; variance for cichlids = 134.6 versus variance for zebrafish = 72.7; F-test, p=0.006). This pattern was also true for latency to enter the top (variance for cichlids = 19,941 versus variance for zebrafish = 10,653; F-test, p=0.004), but not for frequency of entries into the top half (variance for zebrafish = 15.56 vs. variance for cichlids = 15.59; F-test, p=0.996). Phenotypic variance in the novel tank test is represented in Figure 1A-B.

### 3.3 Strong species differences in exploratory behaviors in Lake Malawi cichlids

We next investigated the degree to which phenotypic variance in exploratory behaviors (e.g. see Figure 1) is explained by species differences. Across all three assays, nearly every measured dimension of exploratory behavior differed strongly among species. More detailed results are organized by assay below:

#### Novel tank test

In the novel tank test (Fig. 1A-B), the following standard metrics of exploratory behavior were analyzed across eight species: total time spent in the top half, latency to enter the top half, total number of entries into the top half, and total distance traveled. In addition to these metrics, we also analyzed the average distance from the tank bottom, and the average distance from the tank corners. One-way ANOVAs revealed strong effects of species on total time spent in the top half (F_7,102_=8.64; p=2.74×10^-8^; Eta-squared=0.37, Fig. 2A), latency to enter the top half (F_7,102_=5.44; p=2.50×10^-5^; Eta-squared=0.27), total number of entries into the top half (F_7,102_=8.56; p=3.21×10^-8^; Eta-squared=0.37), total distance traveled (F_7,102_=8.30; p=5.38×10^-8^; Eta-squared=0.36), average distance from the tank bottom (F_7,102_=12.48; p=1.86×10^-11^; Eta-squared=0.46), and average distance from the tank corners (F_7,102_=8.21; p=6.49×10^-8^; Eta-squared=0.36). Pairwise differences between species are shown in Supplementary Figures 1A and 2A-D. Notably, the B/U sand-dweller *Mchenga conophoros* differed strongly from multiple other species in every dimension of behavior analyzed in this test, in every case exhibiting “more exploratory” behavioral phenotypes.

**Figure 2.**
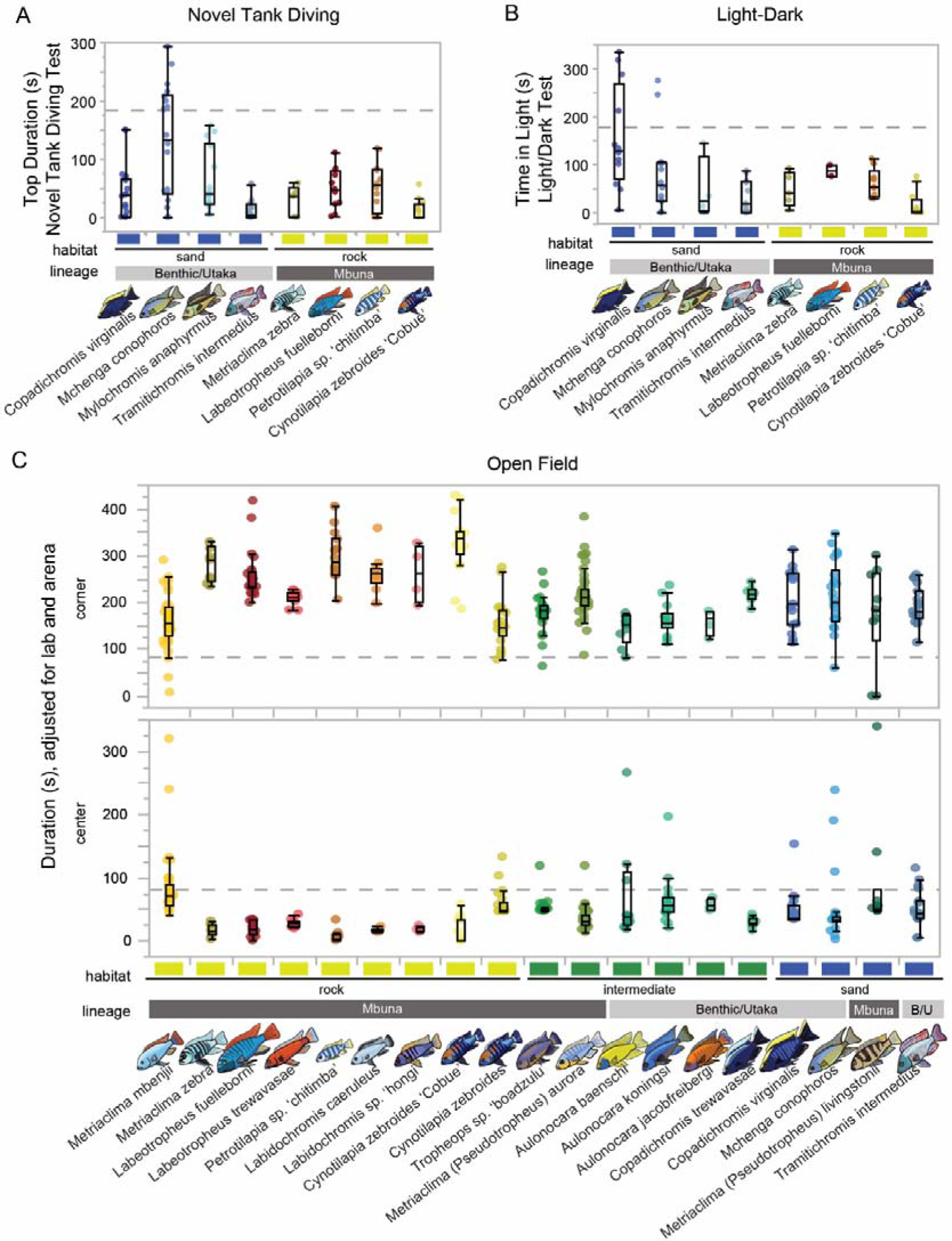
Exploratory behaviors differ strongly between species in Lake Malawi cichlids. Species differences were observed for every measured dimension of exploratory behavior across all three assays. This figure illustrates representative dimensions of behavior in each assay to highlight these differences: time spent in the top half of the novel tank test (A), time spent in the light half of the light-dark test (B), and time spent in the corner and center regions in the open field test (C). For all panels, microhabitat (rock=yellow; intermediate=green; sand=blue) and evolutionary radiation (Mbuna=dark gray; shallow/benthic and utaka=light gray) are color coded and labeled. Dotted lines in all panels indicate null expected values.

#### Light-dark test

For the light-dark test (Fig. 1C-D), total time spent in the light half (Fig. 2B), latency to enter the light half, total number of entries into the light half, and total distance traveled in the light half were analyzed. One-way ANOVAs revealed a significant effect of species on total time spent in the light half (F_7,63_=4.95; p=1.67×10^-4^; Eta-squared=0.35, Fig. 2B), latency to enter the light half (F_7,63_=4.42; p=4.75×10^-4^; Eta-squared=0.33), total number of entries into the light half (F_7,63_=2.54; p=0.023; Eta-squared=0.22), and total distance traveled in the light half (F_7,63_=2.87; p=0.012; Eta-squared=0.24). Pairwise differences between species are shown in Supplementary Figures 1B and 2E-G. Notably, the mbuna rock-dweller *Cynotilapia zebroides ‘Cobue’* exhibited the longest latencies to enter the light half of any species, differing significantly from several other species (Supplementary Figure 2F).

#### Open field test

In the open field test (Fig. 1E-F), time spent in corner regions, corner entries/exits, time spent in the center, center entries/exits, total distance traveled, and speed change over time were analyzed. Because this assay was conducted using two different square arena sizes at two different test locations, the data was analyzed using a one-way ANOVA including an error term with arena size nested within test site. These analyses revealed strong species differences in time spent in the corner regions (F_18,319_=8.928; p<2×10^-16^; Eta-squared=0.33, Fig. 2D top panel), corner entries/exits (F_18,319_=8.901, p<2×10^-16^, Eta-squared=0.33), time spent in the center region (F_18,319_=4.77; p=2.00×10^-9^; Eta-squared=0.21, Fig. 2D bottom panel), center entries/exits (F_18,319_=8.57; p<2×10^-16^; Eta-squared=0.33), total distance traveled (F_18,319_=6.03; p=1.34×10^-12^; Eta-squared=0.25), and speed change over time (F_18,319_=9.20; p<2×10^-16^; Eta-squared=0.34). There were many pairwise differences between species in open field behavior, as shown in Supplementary Figure 1C and 3A-E.

### 3.4 Microhabitat explains variation in open field behaviors

We next investigated whether variation in open field behaviors was associated with microhabitat across 19 species spanning three microhabitats (rock, sand, intermediate). Controlling for variation explained by phylogenetic factors (evolutionary radiation, genus, and species), we found significant associations between microhabitat and multiple open field behaviors. These results are organized into three lines of analysis below (two linear mixed effect regression models, and one MANOVA model; see “Associations among microhabitat, radiation, and open field behaviors” under “Methods” above for full statistical models).

Linear mixed effects regression revealed significant relationships between microhabitat (rock, sand, or intermediate) and open field behavior, while simultaneously accounting for variation explained by phylogenetic factors. Microhabitat (rock, sand, or intermediate) was significantly associated with the number corner entries/exits (F=5.61, p=0.014, Fig. 3A), and this effect was driven by intermediate species entering and exiting the corners more than sand-dwellers (39.4 ± 11.84 more entries, t=3.329, Tukey’s HSD p=0.0096), as well as a trend toward rock-dwelling species entering and exiting the corners more than sand-dwellers (36.2 ± 15.11 more entries, t=2.40, Tukey’s HSD p=0.069). This effect was consistent in direction and statistically significant (Tukey’s p<0.05) regardless of whether test site was included in the model. Microhabitat was also associated with entries/exits to and from the center region (F=12.66, p=5.72×10^-6^, Fig. 3C). The rock microhabitat was associated with more center entries/exits than sand (6.5 ± 2.14 more entries, t=3.04, Tukey’s HSD p=0.0074) and intermediate (4.4 ± 0.94 more entries, t=4.70, Tukey’s HSD p=1.15×10^-5^). The relationship between microhabitat and center entries/exits was not statistically significant or trending when test site was removed from the model (Tukey’s HSD p>0.10 for both effects). A trend was also observed between microhabitat and total distance traveled (F=4.42, p=0.053, Fig. 3E), with intermediate species swimming farther during the test compared to sand-dwellers (1015 ± 358 cm further, t=2.84, Tukey’s HSD p=0.0571), and this effect was consistent in direction and statistically significant when test site was removed from the model (Tukey’s HSD p=0.036). Microhabitat was not significantly associated with time spent in corner regions (F=0.41, p=0.673, Fig. 3B) or time spent in the center region (F=0.70, p=0.512, Fig. 3D), or change in speed over time (F=0.240, p=0.79, Fig. 3F).

**Figure 3.**
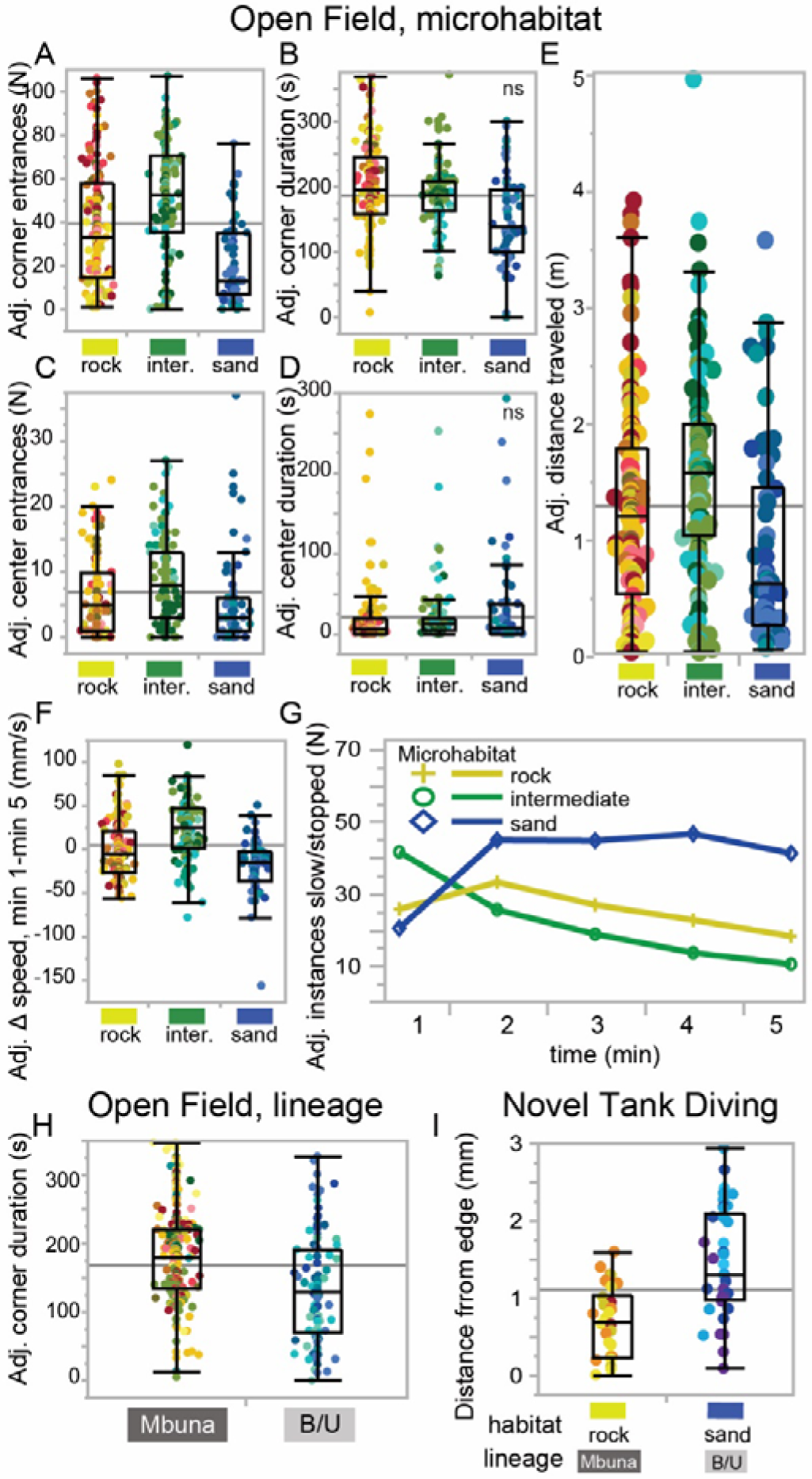
Exploratory behaviors are associated with microhabitat and evolutionary radiation. Variation in open field and novel tank behaviors explained by microhabitat and estimates from linear regression. Controlling for phylogenetic factors, corner entrances/exits (A), but not time spent in the corner (B), are significantly associated with microhabitat (p=0.014). Center entries/exits (C), but not time in the center (D), are also associated with microhabitat (p<0.0001). The association between microhabitat and distance traveled (E) is trending towards significance (p=0.053). Microhabitat is not associated with change in speed over the course of the open field test (F); however, microhabitat is significantly associated with instances of stopping and slowed swimming (G) throughout the open field test (p<0.0001). Exploratory behaviors are also associated with evolutionary radiation (mbuna versus shallow/deep benthic and utaka, B/U). Controlling for variance explained by intermediate versus non-intermediate microhabitat, mbuna species spent significantly more time in corners (H), less time in the center (not shown), and made more entries/exits into the center compared to B/U species (p<0.05 for all). Mbuna rock-dwellers also remained significantly closer to the edges/corners in the novel tank test compared to B/U sand-dwellers (I; p=0.038).

**Figure 3.**
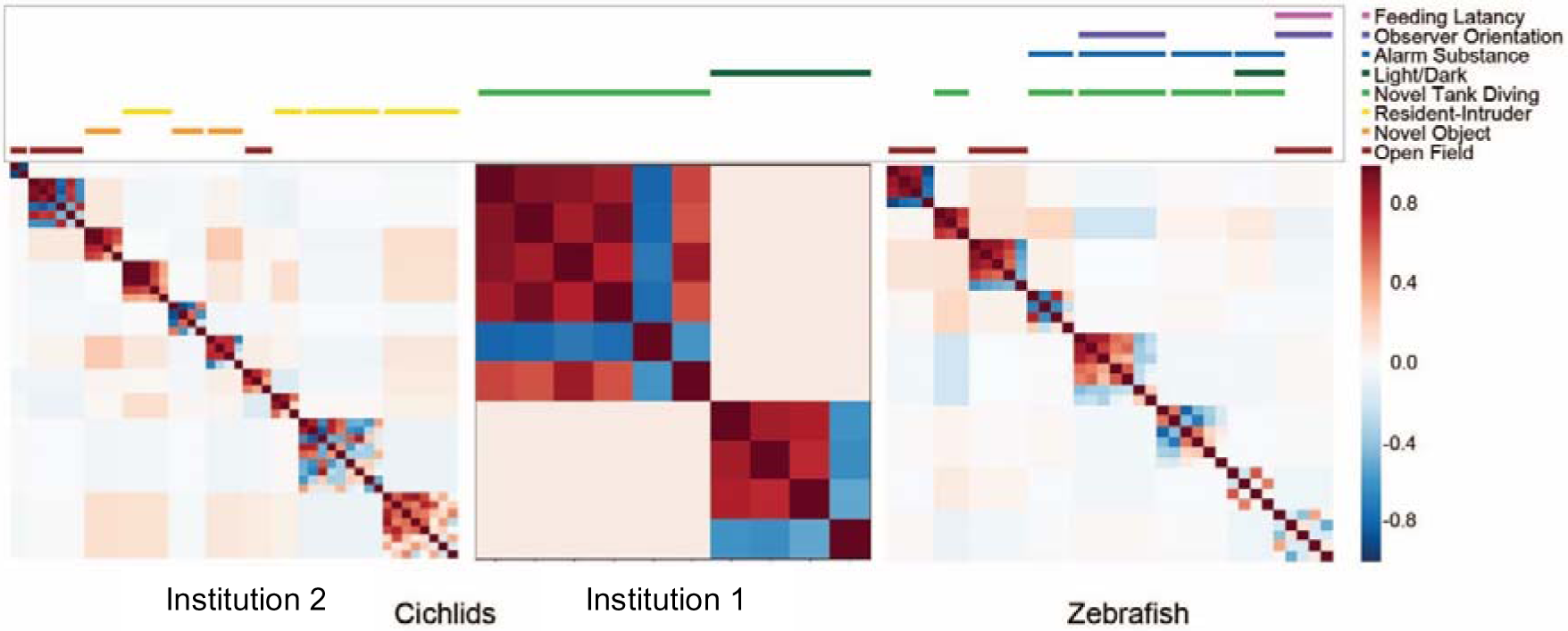
Behavioral modularity analysis across assays in Lake Malawi cichlids and high- and low-exploratory strains of zebrafish. MMC analysis of correlated behaviors across contexts shows extensive clustering within assays in cichlids (INSTITUTION 2 and INSTITUTION 1). In contrast, high- and low-exploratory strains of zebrafish show extensive clustering across assays, indicating strong correlations in behaviors across contexts. Each entry into the matrix is a single behavioral measurement (such as seconds in the corner (open field), or latency to enter the top of the arena (novel tank)). The modules show the pairwise correlations between behavioral measurements across all individuals, with dark red indicating a strong positive correlation and dark blue indicating a strong negative correlation. The color-coded line(s) above each heatmap indicate the behavioral assay(s) represented in each module.

To further investigate the relationships between microhabitat and behavior, we tested a second model in which each microhabitat was designated as either intermediate (rock/sand interface) or non-intermediate (rock or sand). This model allowed effects of microhabitat to be more fully dissociated from effects of evolutionary radiation. The model also included an interaction term to test whether the intermediate microhabitat was differentially associated with behavior between evolutionary radiations. Consistent with our findings from above, this model revealed a strong association between the intermediate microhabitat and entries/exits to and from the corner regions (F=27.08, p=0.0011, Fig. 3A), and this relationship differed between evolutionary radiations (F=6.7945, p=0.041): although intermediate species made more entries/exits to and from the corner regions than non-intermediates in both lineages, the difference was much greater within the B/U radiation (estimated difference of 50.2 ± 10.36 more entries by intermediates vs. non-intermediates, t=2.61, p=0.00056) compared to the mbuna radiation (estimated difference of 17.1 ± 7.57 more entries by intermediates vs. non-intermediates, t=2.26, p=0.10). The model also supported the association between intermediate microhabitat and distance traveled (F=9.17, p=0.018, Fig. 3E), with intermediate species traveling farther than non-intermediates (estimated difference of 729 ± 241 cm farther, t=3.028, p=0.018). Lastly, the model revealed that the intermediate microhabitat was differentially related to swimming speeds in the mbuna versus B/U radiations (F=5.70, p=0.030): controlling for microhabitat, mbuna intermediate species slowed down more than their non-intermediate counterparts during the test (32.1 ± 12.46 mm/s greater decrease in swimming speed, t=2.572, p=0.027), and this pattern was reversed but not statistically significant in B/U species (12.9 ± 15.52 mm/s greater *increase* in swimming speed, t=0.34, p=0.41). All of the above effects were statistically significant (Tukey’s p<0.05) when test site was excluded from the model, with the exception of the interaction between radiation, microhabitat, and change in speed (Tukey’s p>0.10). The full linear regression results for open field behavior, including estimates for pairwise differences between microhabitats, are presented in Tables 1 and 2.

**Table 1.** Associations among microhabitat, evolutionary radiation, and open field behaviors. Summary of linear mixed effect regression output for associations among microhabitat, evolutionary radiation (mbuna versus B/U), and exploratory behaviors in the open field test. The full regression model is shown at the top of the table and was fit to open field behavioral data, with bold italicized terms representing fixed effects, and non-bold italicized terms representing random effects, with nested terms in parentheses. For each behavior, the standard output from linear regression in R is summarized, organized by fixed effect and then by pairwise comparisons between each level of each fixed effect. The output includes the estimated difference between levels of each fixed effect, as well as the standard error, t-statistic, and Tukey’s HSD p-value for each difference. Behaviors, fixed effects, and Tukey’s HSD p-value for each difference. Behaviors, fixed effects, and significant (Tukey’s HSD p<0.05) in the model. Behaviors, fixed effects, and levels of fixed effects that are underlined were found to be statistically significant or trending in this model and also in a second model excluding test site (Tukey’s HSD p<0.10 in both models). Asterisks indicate levels of significance (* for p<0.05; ** for p<0.005; *** for p<0.0005; **** for p<0.00005; and “.” for p<0.10).

| OF behavior ~ *microhabitat + radiation + (genus/species) + test site + arena size* |  |  |  |  |  |  |
| --- | --- | --- | --- | --- | --- | --- |
| Behavior | Fixed effect | Comparison | Estimate | S.E. | t | Tukey's p |
| Time in corners (s) | Microhabitat | Rock-Sand | 41.5 | 32.4 | 1.281 | 0.4237 |
|  |  | Rock-Inter | 26.6 | 20.7 | 1.282 | 0.4249 |
|  |  | Sand-Inter | -14.9 | 24.6 | -0.61 | 0.8189 |
|  | Radiation | Mbuna-Benthic/Utaka | 23.9 | 24.4 | 0.979 | 0.3404 |
| <u>Corner entries and exits</u> | <u>Microhabitat</u> * | <u>Rock-Sand</u> | 36.2 | 15.1 | 2.398 | 0.0689 . |
|  |  | Rock-Inter | -3.2 | 9.3 | -0.344 | 0.9371 |
|  |  | <u>Sand-Inter</u> | **-39.4** | **11.8** | **-3.329** | **0.0096 \*\*** |
|  | Radiation | Mbuna-Benthic/Utaka | -16.4 | 13 | -1.265 | 0.2283 |
| Time in center (s) | Microhabitat | Rock-Sand | -18.2 | 18.8 | -0.966 | 0.6070 |
|  |  | Rock-Inter | -13.4 | 11.9 | -1.125 | 0.5138 |
|  |  | Sand-Inter | 4.8 | 14.4 | 0.332 | 0.9412 |
|  | Radiation | Mbuna-Benthic/Utaka | -3.6 | 14.2 | -0.251 | 0.8046 |
| <u>Center entries and exits</u> | Microhabitat **** | **Rock-Sand** | **6.5** | **2.1** | **3.041** | **0.0074 \*** |
|  |  | **Rock-Inter** | **4.4** | **0.9** | **4.701** | **1.2x10<sup>-5</sup>\*\*\*\*** |
|  |  | Sand-Inter | -2.1 | 2.1 | -1.008 | 0.5724 |
|  | <u>Radiation</u> ** | <u>Mbuna-Benthic/Utaka</u> | **-9.6** | **2.6** | **-3.640** | **0.0029 \*\*** |
| <u>Distance traveled (cm)</u> | <u>Microhabitat</u> . | Rock-Sand | 728 | 452 | 1.611 | 0.2731 |
|  |  | Rock-Inter | -288 | 290 | -0.991 | 0.5959 |
|  |  | <u>Sand-Inter</u> | -1015 | 358 | -2.839 | 0.0572 . |
|  | Radiation | Mbuna-Benthic/Utaka | -308 | 338 | -0.909 | 0.3771 |
| Speed change (mm/s) | Microhabitat | Rock-Sand | -28.9 | 20.1 | -1.441 | 0.3419 |
|  |  | Rock-Inter | -18.3 | 13.0 | -1.406 | 0.3604 |
|  |  | Sand-Inter | 10.6 | 16.9 | 0.630 | 0.8054 |
|  | Radiation | Mbuna-Benthic/Utaka | 27.7 | 15.2 | 1.819 | 0.1005 |

**Table 2.** Relationships among intermediate microhabitat, evolutionary radiation, and open field behaviors. Linear mixed effect regression output for relationships among microhabitat (intermediate versus non-intermediate), evolutionary radiation (mbuna versus B/U), the interaction between microhabitat and radiation, and open field behaviors. For each behavior, the standard output from linear regression in R is summarized, as explained above for Table 1. Behaviors, fixed effects, and levels of fixed effects in bold indicate associations that were found to be statistically significant (Tukey’s HSD p<0.05) in the model. Behaviors, fixed effects, and levels of fixed effects that are underlined were found to be statistically significant or trending in this model and also in a second model excluding test site (Tukey’s HSD p<0.10 in both models). Asterisks indicate levels of significance (* for p<0.05; ** for p<0.005; “.” For p<0.10).

| OF behavior ~ *intermediate + radiation + intermediate\*radiation + (genus/species) + test site + arena size* |  |  |  |  |  |  |
| --- | --- | --- | --- | --- | --- | --- |
| Behavior | Fixed effect | Comparison | Estimate | S.E. | t | Tukey's p |
| <u>Time in corners (s)</u> | Inter | Inter - Non-inter | -1.1 | 17.2 | -0.066 | 0.9481 |
|  | <u>Radiation</u> * | <u>Mbuna - Benthic/Utaka</u> | **45.6** | **17.2** | **2.658** | **0.0175 \*** |
|  | Inter*Radiation | Mbuna (Inter - Non-inter) | -16.9 | 23.7 | -0.711 | 0.8912 |
|  |  | Benthic/Utaka (Inter - Non-inter) | 14.6 | 25.0 | 0.583 | 0.9358 |
| <u>Corner entries and exits</u> | <u>Inter</u> ** | <u>Inter - (Non-inter)</u> | **33.7** | **6.5** | **5.196** | **0.0011 \*\*** |
|  | Radiation | Mbuna - Benthic/Utaka | 4.69 | 10.4 | 0.452 | 0.6650 |
|  | <u>Inter*Radiation</u> * | Mbuna (Inter - Non-inter) | 17.1 | 7.6 | 2.264 | 0.2775 |
|  |  | <u>Benthic/Utaka (Inter - Non-inter)</u> | **50.2** | **10.4** | **4.848** | **0.0027 \*\*** |
| <u>Time in center (s)</u> | Inter | Inter - (Non-inter) | -16.4 | 9.7 | -1.685 | 0.1138 |
|  | <u>Radiation</u> * | <u>Mbuna - Benthic/Utaka</u> | **-24.0** | **10.4** | **-2.306** | **0.0472 \*** |
|  | Inter*Radiation | Mbuna (Inter - Non-inter) | -16.1 | 11.0 | -1.473 | 0.5404 |
|  |  | Benthic/Utaka (Inter - Non-inter) | -16.7 | 15.2 | -1.100 | 0.6941 |
| <u>Center entries and exits</u> | Inter | Inter - (Non-inter) | 2.0 | 1.7 | 1.190 | 0.2531 |
|  | <u>Radiation</u> * | <u>Mbuna - Benthic/Utaka</u> | **-4.52** | **1.9** | **-2.356** | **0.0428 \*** |
|  | Inter*Radiation | Mbuna (Inter - Non-inter) | 0.0 | 2.06 | 0.014 | 1.00 |
|  |  | Benthic/Utaka (Inter - Non-inter) | 4.0 | 2.6 | 1.545 | 0.4366 |
| <u>Distance traveled (cm)</u> | <u>Inter</u> * | <u>Inter - (Non-inter)</u> | **729** | **241** | **3.028** | **0.0183 \*** |
|  | Radiation | Mbuna - Benthic/Utaka | 127 | 230 | 0.552 | 0.5901 |
|  | Inter*Radiation | Mbuna (Inter - Non-inter) | 433 | 315 | 1.376 | 0.5368 |
|  |  | <u>Benthic/Utaka (Inter - Non-inter)</u> | 1024 | 351 | 2.914 | 0.0753 . |
| <u>Speed change (mm/s)</u> | Inter | Inter - (Non-inter) | 9.56 | 10.4 | 0.915 | 0.3718 |
|  | <u>Radiation</u> . | <u>Mbuna - Benthic/Utaka</u> | 19.1 | 10.1 | 1.902 | 0.0863 . |
|  | <u>Inter*Radiation</u> * | Mbuna (Inter - Non-inter) | 32.06 | 12.5 | 2.573 | 0.1060 |
|  |  | Benthic/Utaka (Inter - Non-inter) | -12.94 | 15.5 | -0.834 | 0.8377 |

Microhabitat was also associated with additional patterns of movement over time in the open field test (repeated measures MANOVA, full model F_(4,336)_=11.81, p<0.0001). Both frequency of freezing (F_(2,336)_=15.64 p<0.0001) and the pattern of freezing over time (Wilks’ Lambda value 0.866, approx. F_(8,666)_=6.23, p<0.0001) were associated with microhabitat. Intermediate species initially froze more frequently and exhibited a decrease in slowed swimming as the assay progressed, whereas sand species initially froze less but tended to freeze more as the assay progressed (Fig. 3G).

### 3.5 Open field behaviors differ between mbuna and benthic/utaka evolutionary radiations

The same linear mixed effects regression models described above were used to test for relationships between evolutionary radiation and behavior. Controlling for variance explained by microhabitat, these models revealed that mbuna vs. B/U radiations differed in time spent in corner regions (F=7.065, p=0.018, Table 2, Fig. 3H), time spent in the center region (F=5.32, p=0.047, Table 2), and entries/exits to and from the center region (Model 1, F=13.25, p=0.0029, Table 1; Model 2, F=5.55, p=0.043, Table 2). In comparison to B/U species, mbuna species spent more time in the corner regions (45.6 ± 17.2 seconds, t=2.658, p=0.0175), less time in the center region (24.0 ± 10.4 seconds, t=2.306, p=0.0472), and made fewer entries/exits to and from the center region (4.5 ± 1.9 fewer entries/exits, t=2.356, p=0.0428). The direction of all three of these effects was the same at both test sites, and all three of these effects were consistent in direction and statistically significant or trending towards significance when test site was excluded from the model (Tukey’s p<0.10 for all). A trend toward differences in speed change over time was also observed between radiations (F=3.62, p=0.086), with mbuna species slowing more as the assay progressed compared to B/U species (19.1 ± 10.1 mm/s greater decrease in swimming speed, t=1.902, p=0.0863). This effect was consistent in direction at both test sites and was statistically significant and consistent in direction when test site was excluded from the model (p=0.032). Notably, for all (6/6) open field behaviors analyzed, significant or trending relationships with microhabitat and/or evolutionary radiation were found regardless of whether test site was included in the model.

### 3.6 Novel tank corner behaviors differ between mbuna rock-dwellers and benthic/utaka sand-dwellers

Because of the strong differences between mbuna versus B/U radiations in open field behavior, we also reanalyzed novel tank data, in which four mbuna rock-dwelling species and four B/U sand-dwelling species were tested. Consistent with differences in corner behavior in the open field test, a linear mixed effects regression showed that mbuna rock-dwellers remained significantly closer to outer corner regions compared to B/U sand-dwellers in the novel tank test (0.56 ± 0.23 cm closer, t=2.43; p=0.038, Fig. 3I), but did not differ in the other measured dimensions of novel tank behavior.

### 3.7 Behavioral modularity versus integration: exploratory behaviors are not strongly correlated across contexts in Lake Malawi cichlids

We next investigated evidence for phenotypic integration versus phenotypic modularity of exploratory behaviors in Lake Malawi cichlids. To do this, we analyzed correlations of exploratory behaviors across novel contexts using MMC, which identifies clusters of covariation in large multivariate datasets. We reasoned that if exploratory behaviors are phenotypically integrated, we would expect to observe strong correlations in exploratory behaviors across novel contexts. In contrast, if exploratory behaviors are modular, we would expect to observe weak or no correlations in exploratory behaviors across contexts. As a ground truth and control, we first demonstrated that MMC could reveal clusters of correlated behaviors across contexts by re-analyzing a previously published dataset from selectively bred high- and low-exploratory strains of wild-derived zebrafish. In this study, subjects were phenotyped across a battery of assays (including the novel tank, light-dark, and open field tests among others) and were found to exhibit correlated behaviors across assays (Wong *et al*, 2012). As expected, MMC revealed extensive across-assay clustering in this dataset, with 5/8 (62.5%) clusters spanning multiple assays (including clustering across novel tank and light-dark assays). We then applied MMC to two independent Lake Malawi cichlid datasets, one in which subjects were phenotyped in the novel tank test and light-dark test at INSTITUTION 1, and a second, separate set of experiments in which subjects were phenotyped in a rectangular open field test, resident-intruder test, and novel object test at INSTITUTION 2. For both Malawi cichlid datasets, behavioral clusters grouped exclusively within assay rather than across assays—0/10 modules from the INSTITUTION 2 data set and 0/2 modules from the INSTITUTION 1 data spanned multiple assays.

We phenotyped a wide array of Lake Malawi cichlid species in three classic novel environment assays for the first time. Collectively, Lake Malawi cichlids showed strong behavioral responses that mirrored those of other teleost lineages in all three assays (novel tank test, light-dark test, open field test), spending less time in the top half in the novel tank test, the light half in the light-dark test, and the center region in the open field test (Caio Maximino, de Brito, de Moraes et al., 2007; Stewart, Cachat, Wong et al., 2010; Stewart, Gaikwad, Kyzar et al., 2012; Yoshida, Nagamine, & Uematsu, 2005). The directions of bias in the light-dark and open field tests also match biases displayed by rodents in similarly designed assays: for example, mice and rats spend less time in the light zone in the light-dark test and the center region in the open field test (Bailey & Crawley, 2009; Ramos, Berton, Mormède et al., 1997). Taken together, these results support conserved behavioral and/or stress responses to specific types of novel stimuli that are shared between Lake Malawi cichlids and other teleosts, and more broadly across vertebrates.

Although the direction of these biases was consistent in all species tested, some species exhibited significantly weaker or stronger biases compared to others. For example, the B/U sand-dweller *Mchenga conophoros* spent significantly more time in the top half of the novel tank test compared to every other species tested, and the B/U sand-dweller *Copadichromis virginalis* spent significantly more time in the light half of the light-dark test compared to several other species. To our knowledge, *Copadichromis virginalis* is the only teleost species tested to date that does not exhibit clear scototactic behavioral responses in the light-dark paradigm. Another notable outlier was *Metriaclima mbenjii,* a mbuna rock-dweller, which spent less time in the corner regions of the open field test compared to most other species. Interestingly, *Metriaclima mbenjii* is also atypical in its ecology, being the only *Metriaclima* species found in the sediment-free environment at Mbenji, and (unlike typical rock-dwellers) foraging in large schools in the clear open water (Konings, 2007). Future studies are needed to understand the ecological and/or biological factors contributing to these divergent behavioral phenotypes. The ability to hybridize Lake Malawi cichlids across species boundaries (and between the mbuna and B/U radiations) is a promising strategy for identifying natural genetic variants contributing to behavioral divergence.

We also investigated the degree of phenotypic variance in exploratory behaviors in Lake Malawi cichlids. To place our analyses in a frame of reference, we measured phenotypic variance in novel tank behaviors among Lake Malawi cichlids and among three laboratory strains of zebrafish that were tested with the same parameters in a previous study: two wild-derived strains that were selectively bred for extreme and opposite exploratory behavioral phenotypes, and a common wild-type laboratory strain (AB). Notably, the average genetic divergence between Lake Malawi cichlid species is less than between common laboratory strains of zebrafish (Loh, Katz, Mims et al., 2008). We found that Lake Malawi cichlids collectively exhibited significantly greater variance in multiple dimensions of exploratory behavior compared to the zebrafish strains, including time spent in the top half and entries into the top half. These results suggest that natural evolution in Lake Malawi cichlids has resulted in extreme phenotypic diversity in exploratory behaviors, similar to other complex traits such as morphology and color patterning.

We next tested the extent to which this phenotypic diversity is explained by species differences. Strong species differences were observed for nearly every dimension of exploratory behavior analyzed across all assays. Taken together, these results show that the extreme diversity in exploratory behaviors in Lake Malawi cichlids is partially explained by rapid and strong behavioral divergence along species lines. This is consistent with findings in other vertebrate lineages, in which behavioral responses to novel stimuli have rapidly diverged between closely-related species of birds and mammals (Cowan, 1977; Greenberg, 2003; C. Mettke-Hofmann, Winkler, Hamel et al., 2013; Claudia Mettke-Hofmann, Winkler, & Leisler, 2002). Considering the low genetic divergence and ability to hybridize between species, these results further demonstrate that Lake Malawi cichlids are a powerful complementary system to traditional laboratory models for understanding the genetic basis of naturally evolved species differences in exploratory behaviors.

We investigated the ecological basis of species differences in exploratory behavior by phenotyping 19 species spanning three Lake Malawi microhabitats (rock, sand, and intermediate) in the open field test. Controlling for variation explained by phylogenetic factors, microhabitat was associated with multiple dimensions of open field behavior, including entries/exits to and from the corners, entries/exits to and from the center, and total distance traveled. Notably, intermediate species traveled significantly farther and made significantly more entries/exits to and from the corners compared to non-intermediate species. It is difficult to speculate how these differences may translate to behavior in nature; however, these data suggest that habitat divergence—from the canonical rock and sand ecotypes into the intermediate zone—is associated with behavioral divergence in Lake Malawi cichlids. Future work is needed to understand the causal relationships between behavioral divergence, invasion of the intermediate habitat, and adaptation to ecological conditions in the intermediate habitat. Interestingly, relationships between the intermediate microhabitat and behavior differed between the mbuna and B/U radiations for multiple dimensions of open field behavior, including corner entries/exits and speed change over time. These results support the idea that lineage-specific behavioral specializations are associated with divergence into the intermediate habitat. The mbuna versus B/U radiations differ both genomically and ecologically (with mbuna species tending to inhabit rock habitats, and B/U species tending to inhabit sand). Therefore, it is unclear whether these behavioral differences have been driven by divergence between the underlying genomic backgrounds of these radiations, and/or by differences in the ecological transitions from rock to intermediate zone, versus sand to intermediate zone.

We also investigated whether a major evolutionary radiation in Malawi cichlids (mbuna versus B/U) explains natural variation in exploratory behaviors. Controlling for variation explained by microhabitat, multiple dimensions of open field behavior differed significantly between the mbuna and B/U radiations, including time spent in the corners, time spent in the center, and center entries/exits. In all three cases, the mbuna species exhibited phenotypes that were less exploratory compared to B/U species. Consistent with this pattern, mbuna rock-dwellers also remained significantly closer to the corner regions in the novel tank test compared to B/U sand-dwellers. Taken together, these results provide evidence for divergence in general exploratory phenotypes between two major cichlid radiations in Lake Malawi. One potential explanation for these patterns is that behavioral preferences for edges or corners helps mediate behavioral preferences for the narrow crevasses and caves characteristic of rocky habitats; inversely, a reduced aversion toward open environments may facilitate preferences for and/or invasion of new and potentially more exposed habitats, such as the sandy microhabitats in Lake Malawi. An alternative explanation is that differences in the genomic backgrounds, or random genetic drift between radiations has produced random phenotypic drift in exploratory behaviors. Notably, mbuna versus B/U lineages are separated by known fixed genetic differences as well as neurogenetic and neuroanatomical specializations (e.g. volume of the cerebellum and telencephalon) (Huber, van Staaden, Kaufman et al., 1997; Sylvester, Rich, Loh et al., 2010), highlighting potential targets for future studies of behavioral divergence.

Comparative studies in Lake Malawi cichlids have previously demonstrated modular patterns of covariation for complex traits that are thought to have played a central role in cichlid diversification, including oral jaw morphology and color patterning (R. Craig Albertson, Powder, Hu et al., 2014; Parsons, Cooper, & Albertson, 2011). Similarly, we reasoned that, because behavioral responses to novel stimuli are measurable traits, then correlations of behavioral responses to different novel stimuli can provide evidence for behavioral integration versus behavioral modularity. Following this logic, we tracked individuals across multiple novel contexts to investigate whether patterns of covariation among behavioral responses are modular or integrated. To do this, we applied MMC, a statistical approach designed to identify clusters of covariation in large multivariate datasets. We first applied MMC to a previously published dataset in which laboratory strains of zebrafish were found to exhibit correlated, or syndromic, behaviors across contexts (Baker, Goodman, Santo et al., 2018; Ryan Y Wong, Perrin, Oxendine et al., 2012). In line with previous results, MMC revealed extensive clustering across assays, indicating that behaviors were correlated across contexts, and consistent with integration of behavioral responses in different novel contexts. We then applied MMC to two independent Malawi cichlid datasets, in which subjects were phenotyped in different assays at two separate institutions. In both datasets, behaviors clustered exclusively within assay, consistent with modularity of behavioral responses across novel contexts. Taken together, these results support the idea that, like other complex traits, Lake Malawi cichlids exhibit modular patterns of behavioral variation. Future studies are needed to investigate whether exploratory behaviors are more evolvable in this species assemblage, and whether they have played a causal role in cichlid diversification.

There are several limitations to these experiments. First, these assays do not reflect environmental conditions in Lake Malawi, and therefore it is unclear how behavioral phenotypes in these experiments map onto behavior in natural environments. Additionally, although the number of species investigated was larger than most comparative behavioral investigations, larger samples of species and individuals may uncover additional links between more specific dimensions of ecology and behavioral variation. For example, factors such as diet, resource distribution, population density, developmental stage, turbidity, depth, and/or predation risk may explain species differences in behavioral responses to novel stimuli. Additional factors may also influence behavioral responses to novel stimuli across species, such as developmental stage, sex, or social context. Lastly, our analyses of behavioral modularity versus integration only spanned five contexts and two relatively small subsets of species. Future studies are needed to more systematically investigate behavioral modularity versus integration in across more species and contexts.

Despite these limitations, these experiments constitute a large comparative investigation of exploratory behavioral variation in a previously untested vertebrate system. We phenotype a total of 20 new species in a variety of classic behavioral assays and show conserved behavioral responses that mirror other teleosts and rodents. We demonstrate high phenotypic variance in exploratory behaviors that segregates along species lines. We further show that natural variation in exploratory behaviors are explained by microhabitat and by major evolutionary radiations in Lake Malawi. Lastly, we provide evidence for behavioral modularity in Lake Malawi cichlids. Taken together, these findings provide new insights into the ecology and evolution of exploratory behaviors, and demonstrate Lake Malawi cichlids as a powerful complement to traditional models for investigating the ecological, genetic, and neural factors underlying natural behavioral diversity.

## Acknowledgements

Preparation of this manuscript was supported by an Arnold and Mabel Beckman Foundation Beckman Young Investigator Award to RBR and by NIH grant R01GM101095 to JTS

## Supplementary Methods and Results

### S.1 Effects of test site

Test site was included as a covariate in our linear mixed-effects models, and thus variance in behavioral data explained by test site was controlled for in our primary analyses. Nonetheless, we performed additional analyses to investigate the possibility that effects of test site, or combining the data across institutions, may have influenced our results. We first analyzed the relationships between test site and behavior for the only species that was housed and tested at both sites, *Labeotropheus fuelleborni* (INSTITUTION 1, n=16; INSTITUTION 2, n=7). Controlling for arena size, linear regression showed that test site was not significantly associated or trending with any of the six analyzed open field behaviors: corner time (t=1.33, p=0.20), corner entries/exits (t=0.15, p=0.88), center time (t=0.56, p=0.58), center entries/exits (t=0.86, p=0.39), distance traveled (t=0.54, p=0.60), and speed change (t=1.56, p=0.14).

To further investigate potential effects of test site on our results, we conducted all open field analyses with and without test site as a covariate, and found that the vast majority (11/14, ∼79%) of significant or trending relationships in our primary analyses were also significant or trending when test site was excluded from the models. The few results that changed from statistically significant to p>0.10, as well as all results that were significant or trending in both models, are indicated in Tables 1 and 2.

Although the above analyses suggested that our signals were not driven by differences between test sites, we further addressed this concern by validating that these microhabitat-behavior and radiation-behavior signals were also present when the data was analyzed within test site only. We split our dataset into two “within institution” subsets and used linear mixed effects models to analyze relationships among microhabitat, evolutionary radiation, and open field behaviors within each site independently. By splitting the data, we lost a substantial amount of statistical power, and therefore we used the following simplified models to test these relationships:

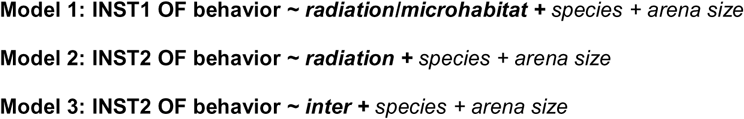

We used a single model with an interchangeable microhabitat/radiation fixed effect term to test INSTITUTION 1 data, because microhabitat could not be separated from radiation in this dataset (Model 1). For the INSTITUTION 2 dataset, we used two models, one to test the relationship between the intermediate microhabitat and open field behaviors (Model 2), and a second to test the relationship between evolutionary radiation and open field behaviors (Model 3). Each model was used to analyze relationships for all six open field behaviors, and thus collectively they resulted in 18 p-values. As in our primary analyses, these analyses resulted in a highly leftward skewed distribution of p-values, with 8/18 relationships being statistically significant (4/18, p<0.05) or trending (4/18, 0.05<p<0.10), ∼4x more than expected by chance (Supplementary Figure 4). In the vast majority of these cases (7/8, ∼88%), the microhabitat-behavior and/or radiation-behavior relationship was matched to a statistically significant (p<0.05) relationship in our primary analyses of the combined data. In the single exception, a trend was observed between mbuna versus B/U species for distance traveled within INSTITUTION 2 (p=0.070), but this relationship was not statistically significant or trending in the combined dataset (Table 1). Taken together, these data strongly suggest that our results were not driven by differences between test sites or by combining the data across institutions.

### S.2 Multiple hypothesis testing: microhabitat, radiation, and open field behaviors

Collectively, our primary analyses tested the relationships between multiple factors (microhabitat, evolutionary radiation, and the interaction between microhabitat and evolutionary radiation) and multiple open field behaviors (time spent in the corners, entries into the corners, time spent in the center, entries into the center, total distance traveled, and change in speed over time). These tests resulted in a total of 30 comparisons and p-values. Although Tukey’s HSD method controlled for multiple comparisons within each level of open field behavior, we also assessed the risk that all comparisons across behavioral levels and linear models generated false positive signals. We used the qvalue Bioconductor package in R to estimate the total proportion of true null findings (π_0_) and the q-value associated with each p-value.

At α=0.05, we would expect to observe approximately ∼1.5 significant results among 30 comparisons, and ∼3 significant results at α=0.10. In contrast, we observed 10 relationships that were significant at the α=0.05 level (∼7x greater than expected by chance), and 12 results that were significant (or “trending”) at α=0.10 (4x greater than expected by chance; Supplementary Figure 4A). To further assess the risk of false positives generated through multiple hypothesis testing, we used our distribution of p-values to estimate the total number of truly null results (π_0_*30), and then calculated the associated q-values for each comparison. Briefly, the q-value for a test is equal to the minimum FDR at which that test would be called significant (Storey, 2002, 2003). Based on our distribution of p-values (see Supplementary Figure 4A), we estimated that ∼18/30 results were likely to be truly null, while ∼12/30 were likely to be true relationships (π_0_=0.569). This estimate fell in line with the number of relationships observed to be significant or trending towards significance (p<0.05 for 10/30 relationships; p<0.10 for 12/30 relationships), and 11/12 (∼92%) of these relationships were matched with q-values below 0.10. Taken together, these results indicate that the vast majority of significant relationships among microhabitat, evolutionary radiation, and exploratory behaviors identified by our models are likely to be true positives.

### S.3 MMC Behavioral Assays

MMC was performed on a set of three behavioral assays that were conducted at INSTITUTION 2 as part of a separate set of experiments. In these experiments, 70 subjects representing five species (*Metriaclima/Pseudotropheus aurora, Metriaclima callainos, Metriaclima lombardoi, Metriaclima pyrsonotos, and Aulonocara baenschi*; n=14 per species; see Supplementary Table 5) were tracked across a resident-intruder test, a novel object test, and a rectangular open field test. The testing parameters for each assay are described below.

#### Rectangular open field assay

The rectangular open field assay was conducted at INSTITUTION 2 as part of a separate study, and these data were analyzed in MMC for the present study. Subjects from five species (n=14 per species, see Supplementary Table 5) were netted from home tanks and transported to the test arena with minimal handling. Each subject was introduced into the center of a 76.0 cm x 46.0 cm rectangular arena filled with 6 cm of water from the common laboratory recirculating system for five minutes (modified from (Godwin et al. 2012)). An overhead digital video camera recorded the position of the fish in the arena over the course of the assay. C-trax v0.5.4 (Branson et al. 2009) generated X, Y coordinates for the position of the center of the fish for a minimum of 10 frames per second over the course of the assay.

#### Resident-intruder test

Individual subjects were measured in the resident-intruder assay as part of a separate study on cichlid aggression (Emily C. Moore & Roberts, 2019). Briefly, following the rectangular open field test, subjects were introduced and acclimated to a 38-liter (51 cm x 28 cm x 33 cm) aquarium containing a terracotta flowerpot territory for three days. After the acclimation period, an age- and size-matched *Labeotropheus trewavasae* subject was introduced as the intruder, and behavior was recorded for five minutes. Additional details on the testing parameters and analysis for this assay are described in Moore and Roberts 2019 (Emily C. Moore & Roberts, 2019).

#### Novel object test

One day after the resident-intruder test, each subject was measured in the novel object test. During this test, a camera was placed overhead, and water and air flow was stopped five minutes prior to the beginning of the test to enable clear video recording and to allow time for subjects to habituate to the change. A snail shell from Lake Malawi was then introduced into the home aquarium and behavior was recorded for 30 minutes with a digital video camera. The position of the most rostral aspect of the head was scored with Manual Tracking plug-in (Cordelieres 2005) for ImageJ (Schneider et al. 2012) in 0.2 second intervals (5 frames per second). Aquarium positioning prevented the entire arena from being filmed, so position analysis was restricted to the front-most 25.4 cm x 26 cm of the tank for all subjects. For the novel object test, total time spent stationary, approaching, and retreating from the object; distance from the object; and approach velocity, retreat velocity, average velocity, and change in velocity over the course of the assay were analyzed.

## Supplementary Tables and Figures

**Supplementary Table 1.** Novel tank place bias between bottom and top regions by species. Each row corresponds to the species labeled in the left column. The following are presented for each species: microhabitat designation, evolutionary radiation (mbuna, Mbu; shallow/deep benthic and Utaka, B/U), test site (Inst.; INSTITUTION 1 vs. INSTITUTION 2), sample size, mean time in bottom zone, mean time in top zone, standard error for time spent in both zones, and two-tailed paired t-test p-values for the difference in time spent between the two zones.

| Species | Microhabitat | Rad | Inst | N | Bottom time (seconds) | Top time (seconds) | S.E. | P<sub>top vs. bottom</sub> |
| --- | --- | --- | --- | --- | --- | --- | --- | --- |
| *Metriaclicma zebra* | Rock | Mbu | 1 | 5 | 332.46 | 27.54 | 13.15 | 2.05x10<sup>-4</sup> *** |
| *Labeothropheus fuelleborni* | Rock | Mbu | 1 | 12 | 310.40 | 49.60 | 10.11 | 3.50x10<sup>-8</sup> *** |
| *Petrotilapia sp. 'chitimba'* | Rock | Mbu | 1 | 12 | 306.82 | 53.18 | 12.66 | 4.71x10<sup>-7</sup> *** |
| *Cynotilapia zebroides 'Cobue'* | Rock | Mbu | 1 | 13 | 348.23 | 11.77 | 4.99 | 1.32x10<sup>-12</sup> *** |
| *Copadichromis virginalis* | Sand | B/U | 1 | 18 | 316.60 | 43.40 | 11.29 | 5.75x10<sup>-10</sup> *** |
| *Mchenga conophoros* | Sand | B/U | 1 | 18 | 229.88 | 130.12 | 23.35 | 0.0421 * |
| *Mylochromis anaphyrmus* | Sand | B/U | 1 | 14 | 295.95 | 64.05 | 15.30 | 2.70x10<sup>-6</sup> *** |
| *Tramitichromis intermedius* | Sand | B/U | 1 | 18 | 346.99 | 13.01 | 4.46 | 5.47x10<sup>-18</sup> *** |

**Supplementary Table 2.** Place bias between light and dark halves of the light-dark test by species. Each row corresponds to the species labeled in the left column. The following are presented for each species: sample size, microhabitat designation, evolutionary radiation (mbuna, Mbu; shallow/deep benthic and Utaka, B/U), mean time in dark zone, mean time in light zone, standard error for time spent in both zones, and two-tailed paired p-value for the difference in time spent between the two zones.

| Species | Microhabitat | Rad | Inst | N | Dark time (seconds) | Light time (seconds) | S.E. | P<sub>light vs. dark</sub> |
| --- | --- | --- | --- | --- | --- | --- | --- | --- |
| *Metriaclima zebra* | Rock | Mbu | 1 | 5 | 312.67 | 47.33 | 17.67 | 0.0011 ** |
| *Labeothropheus fuelleborni* | Rock | Mbu | 1 | 2 | 272.91 | 87.09 | 16.19 | 0.078 |
| *Petrotilapia* sp. 'chitimba' | Rock | Mbu | 1 | 13 | 297.82 | 62.18 | 8.34 | 4.87x10<sup>-9</sup> *** |
| *Cynotilapia zebroides* 'Cobue' | Rock | Mbu | 1 | 12 | 344.07 | 15.93 | 8.15 | 3.12x10<sup>-10</sup> *** |
| *Copadichromis virginalis* | Sand | B/U | 1 | 12 | 203.77 | 156.23 | 32.73 | 0.46 |
| *Mchenga conophoros* | Sand | B/U | 1 | 12 | 275.42 | 84.58 | 27.02 | 0.0036 ** |
| *Mylochromis anaphyrmus* | Sand | B/U | 1 | 4 | 311.83 | 48.17 | 38.02 | 0.028 * |
| *Tramitichromis intermedius* | Sand | B/U | 1 | 11 | 325.74 | 34.26 | 11.07 | 7.74x10<sup>-8</sup> *** |

**Supplementary Table 3.** Place bias between central and peripheral regions of the open field test by species. Each row corresponds to the species labeled in the left column. The following are presented for each species: sample size, microhabitat designation, estimate for mean time in center, estimate for mean time in periphery, standard error for time spent in center and periphery, and two-tailed paired p-value for the difference in time spent in central versus peripheral regions.

| Species | Microhab | Rad | Inst | N | MD | LG | Center time (seconds) | Periphery time (seconds) | S.E. | P<sub>center vs. periphery</sub> |
| --- | --- | --- | --- | --- | --- | --- | --- | --- | --- | --- |
| *Metriaclima mbenjii* | Rock | Mbu | 2 | 39 | 9 | 30 | 62.06 | 257.94 | 10.13 | $7.89 \times 10^{-12}$ *** |
| *Metriaclima zebra* | Rock | Mbu | 1 | 9 | 9 | 0 | 17.32 | 302.68 | 3.31 | $5.83 \times 10^{-11}$ *** |
| *Labeotropheus fuelleborni* | Rock | Mbu | 1,2 | 23 | 15 | 8 | 9.27 | 310.73 | 2.13 | $1.10 \times 10^{-27}$ *** |
| *Labeotropheus trewasae* | Rock | Mbu | 2 | 11 | 11 | 0 | 28.09 | 291.91 | 2.01 | $1.04 \times 10^{-14}$ *** |
| *Petrotilapia* sp. 'chitimba' | Rock | Mbu | 1 | 14 | 12 | 2 | 2.56 | 317.44 | 1.09 | $1.91 \times 10^{-22}$ *** |
| *Labidochromis caeruleus* | Rock | Mbu | 2 | 10 | 10 | 0 | 17.17 | 302.83 | 0.80 | $1.75 \times 10^{-17}$ *** |
| *Labidochromis* sp. 'hong'i' | Rock | Mbu | 2 | 4 | 4 | 0 | 19.37 | 300.63 | 2.50 | $8.08 \times 10^{-6}$ *** |
| *Cynotilapia zebroides* | Rock | Mbu | 2 | 21 | 0 | 21 | 26.22 | 293.78 | 4.94 | $1.92 \times 10^{-17}$ *** |
| *Cynotilapia zebroides* 'Cobue' | Rock | Mbu | 1 | 18 | 9 | 9 | 4.90 | 315.10 | 3.30 | $1.20 \times 10^{-19}$ *** |
| *Tropheops* sp. 'Boadzulu' | Intermediate | Mbu | 2 | 29 | 0 | 29 | 20.29 | 299.71 | 2.48 | $1.37 \times 10^{-30}$ *** |
| *Metriaclima* (*Pseudotropheus*) *aurora* | Intermediate | Mbu | 2 | 55 | 48 | 7 | 29.94 | 290.06 | 2.20 | $5.78 \times 10^{-51}$ *** |
| *Aulonocara baenschii* | Intermediate | B/U | 2 | 9 | 9 | 0 | 73.77 | 246.23 | 28.32 | 0.0121 * |
| *Aulonocara koningsi* | Intermediate | B/U | 2 | 18 | 10 | 8 | 47.91 | 272.09 | 10.50 | $3.87 \times 10^{-9}$ *** |
| *Aulonocara jacobfreibergi* | Intermediate | B/U | 2 | 4 | 0 | 4 | 36.52 | 283.48 | 13.98 | 0.00201 ** |
| *Copadichromis trewasae* | Intermediate | B/U | 2 | 11 | 11 | 0 | 29.75 | 290.25 | 2.23 | $3.24 \times 10^{-14}$ *** |
| *Copadichromis virginalis* | Sand | B/U | 1 | 15 | 0 | 15 | 16.99 | 303.01 | 8.23 | $4.52 \times 10^{-11}$ *** |
| *Mchenga conophoros* | Sand | B/U | 1 | 22 | 10 | 12 | 30.84 | 289.16 | 14.11 | $5.99 \times 10^{-9}$ *** |
| *Tramitichromis intermedius* | Sand | B/U | 1 | 19 | 9 | 10 | 35.07 | 284.93 | 8.03 | $4.39 \times 10^{-12}$ *** |
| *Metriaclima* (*Pseudotropheus*) *livingstonii* | Sand | Mbu | 2 | 10 | 0 | 10 | 56.48 | 263.52 | 30.74 | 0.00622 *** |

**Supplementary Figure 1.**
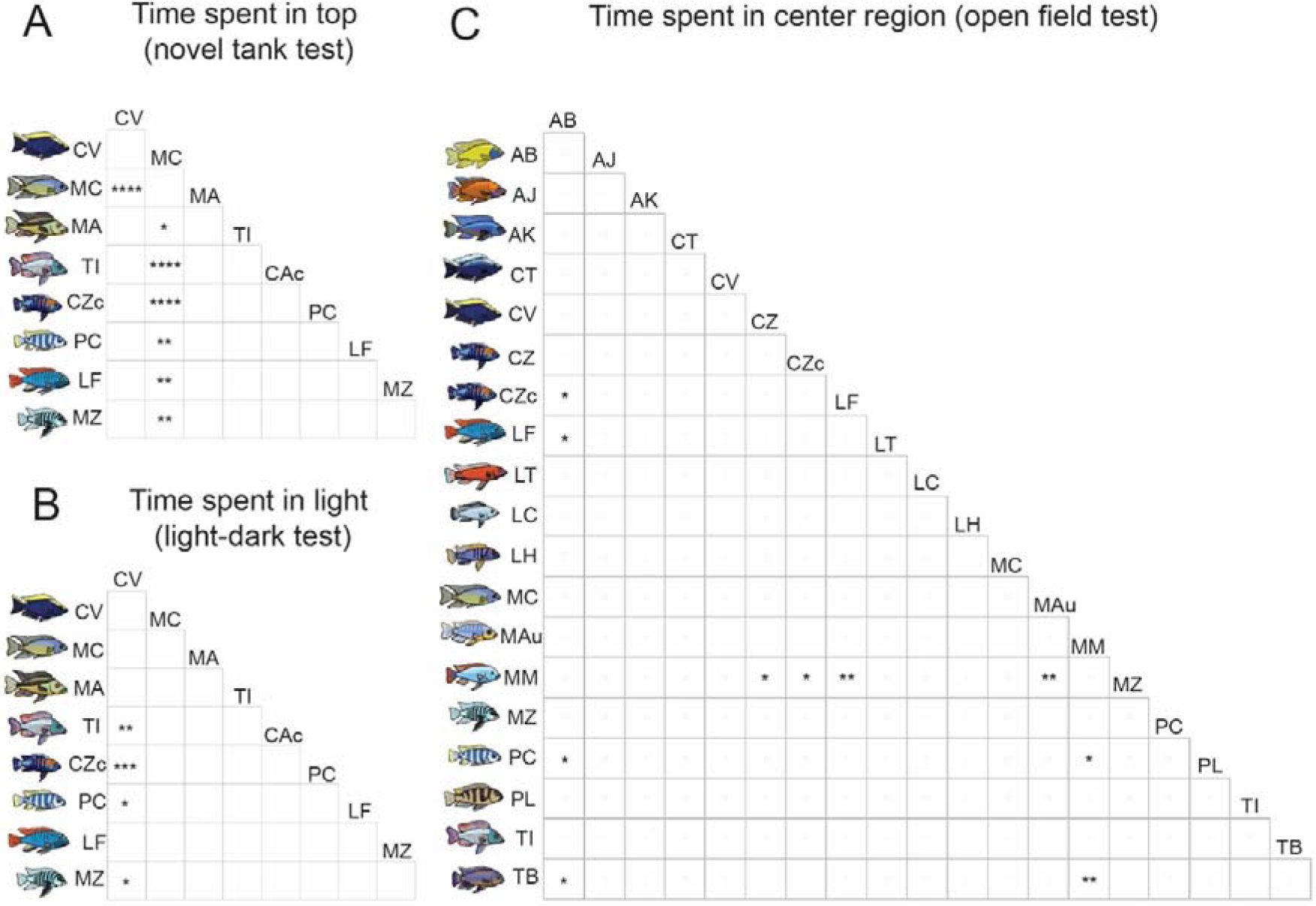
Pairwise species differences in strength of zone preferences across assays. Species differences were present in the amount of time spent in the top half of the novel tank test (A), light half in the light-dark test (B), and center region in the open field test (C). Asterisks indicate levels of significance for post-hoc Tukey’s HSD tests of the difference between species (* p<0.05, ** p<0.005, *** p<0.0005, ****p<5×10^-5^). Species abbreviations are as follows: AB (*Aulonocara baenschii*), AJ (*Aulonocara jacobfreibergi*), AK (*Aulonocara koningsi*), CT (*Copadichromis trewavasae*), CV (*Copadichromis virginalis*), CZ (*Cynotilapia zebroides*), CZc (*Cynotilapia zebroides ‘Cobue’)*, LF (*Labeotropheus fuelleborni*), LT (*Labeotropheus trewavasae*), LC (*Labidochromis caeruleus*), LH (*Labidochromis* sp. ‘hongi’), MC (*Mchenga conophoros*), MA (*Mylochromis anaphyrmus*), Mau (*Metriaclima aurora*), MM (*Metriaclima mbenjii*), MZ (*Metriaclima zebra*), PC (*Petrotilapia sp. ‘chitimba’*), PL (*Pseudotropheus livingstonii*), TI (*Tramitichromis intermedius*), TB (*Tropheops sp. ‘Boadzulu’*).

**Supplementary Figure 2.**
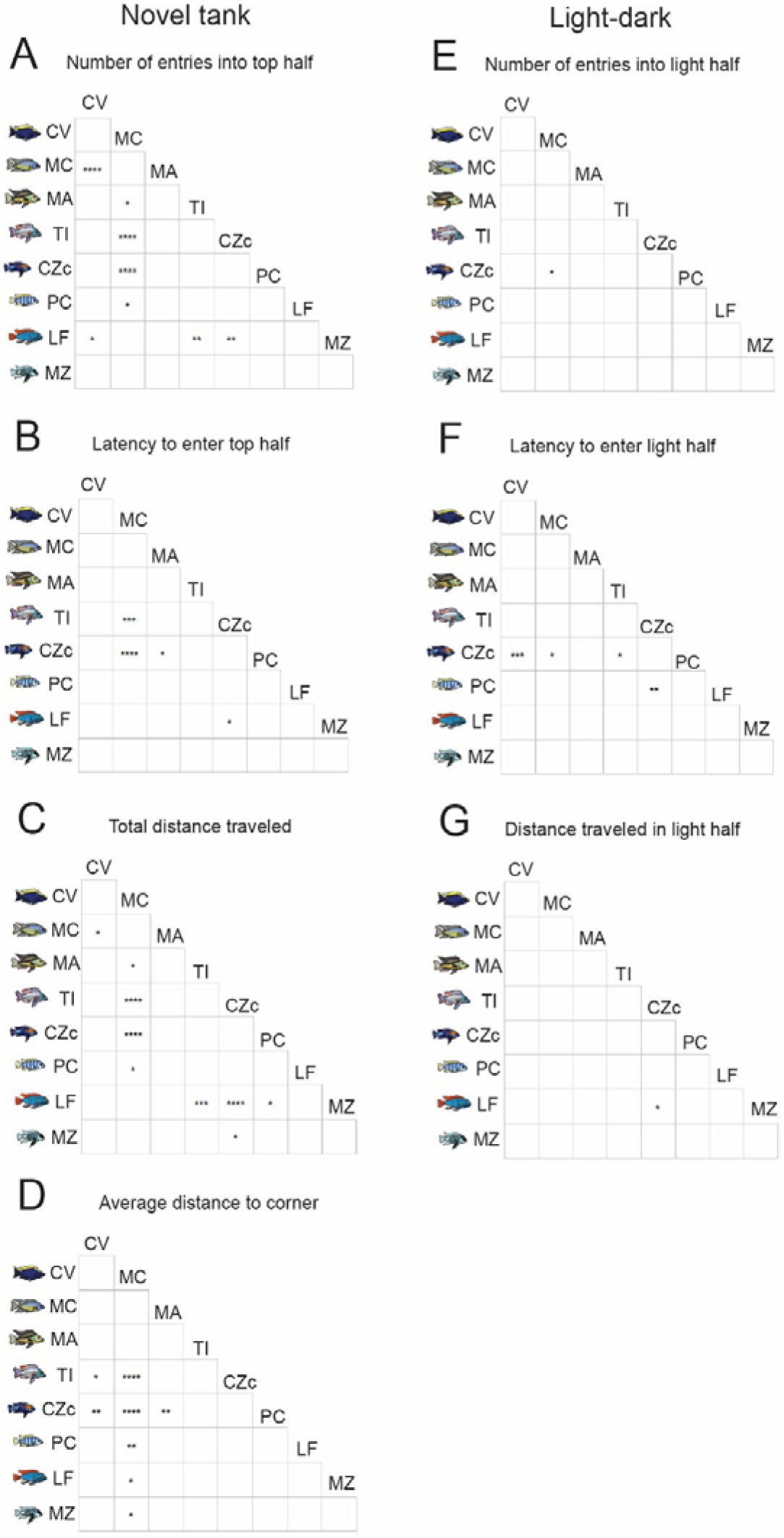
Pairwise species differences in behavior across assays, including the novel tank (A-D) and light-dark tests. Asterisks indicate levels of significance for post-hoc Tukey’s HSD tests of the pairwise differences between species (* p<0.05, ** p<0.005, *** p<0.0005, ****p<5×10^-5^).

**Supplementary Figure 3.**
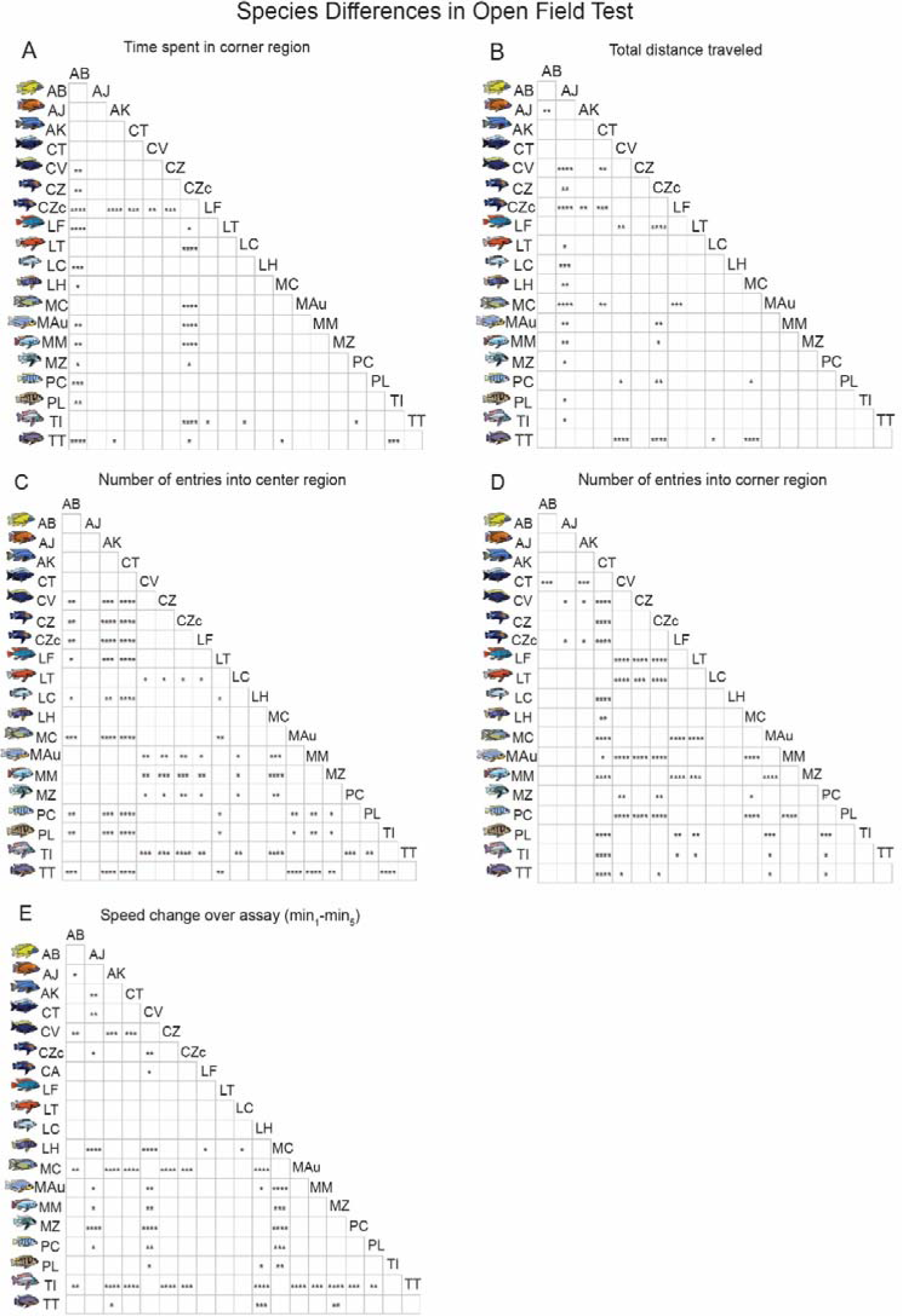
Pairwise species differences in open field behaviors (A-E). Asterisks indicate levels of significance for post-hoc Tukey’s HSD tests of the difference between species (* p<0.05, ** p<0.005, *** p<0.0005, ****p<5×10^-5^).

**Supplementary Table 4.** Samples analyzed for Modularity Modular Clustering analysis, organized by species, institution, sample size, age, and assay. For each species, each “X” indicates the assays in which all subjects sampled were tested. Two independent datasets were analyzed, the first dataset is represented by the first eight species listed, and the second dataset is represented by the last five species listed.

| Species | Institution | N | Age (days) | Novel tank | Light-dark | Resident intruder | Novel object | Open field |
| --- | --- | --- | --- | --- | --- | --- | --- | --- |
| *Copadichromis virginalis* | 1 | 11 | 90-180 | X | X |  |  |  |
| *Cynotilapia zebroides* 'Cobue' | 1 | 11 | 90-180 | X | X |  |  |  |
| *Labeotropheus fuelleborni* | 1 | 2 | 90-180 | X | X |  |  |  |
| *Mchenga conophoros* | 1 | 12 | 90-180 | X | X |  |  |  |
| *Metriaclima zebra* | 1 | 5 | 90-180 | X | X |  |  |  |
| *Mylochromis anaphyrmus* | 1 | 4 | 90-180 | X | X |  |  |  |
| *Petrotilapia* sp. 'chitimba' | 1 | 12 | 90-180 | X | X |  |  |  |
| *Tramitichromis intermedius* | 1 | 10 | 90-180 | X | X |  |  |  |
| *Aulonocara baenschi* | 2 | 14 | >180 |  |  | X | X | X |
| *Metriaclima (Pseudotropheus) aurora* | 2 | 14 | >180 |  |  | X | X | X |
| *Metriaclima callainos* | 2 | 14 | >180 |  |  | X | X | X |
| *Metriaclima lombardoi* | 2 | 14 | >180 |  |  | X | X | X |
| *Metriaclima pyrrsonotos* | 2 | 14 | >180 |  |  | X | X | X |

**Supplementary Table 5.** Total subject counts by species, microhabitat (Rock, Sand, Intermediate=Inter), evolutionary radiation (Mbuna=Mbu, shallow/deep benthic and utaka=B/U), institution (INSTITUTION 1 vs. INSTITUTION 2), and assay (novel tank=NT, light-dark=LD, novel object=NO, open field=OF, resident intruder=RI). Subjects that were tracked in multiple assays are indicated by parentheses, and are only counted once. Numbers that are italicized and marked by asterisks within parentheses indicate individuals that were subjected to three assays (a rectangular open field test, a resident-intruder test, and a novel object test) as part of a separate study. Because these subjects were tracked across multiple assays, their behavior for all three assays was analyzed in MMC. One species, *Labeotropheus fuelleborni,* was housed at both institutions, and is represented in two separate rows to show information about subjects at each institution (indicated as a duplicate species by parentheses).

| Species | Microhab | Rad | Inst | NT | LD | NO | OF | RI | Total |
| --- | --- | --- | --- | --- | --- | --- | --- | --- | --- |
| *Metriaclima mbenjii* | Rock | Mbu | 2 | 0 | 0 | 0 | 39 | 0 | 39 |
| *Metriaclima zebra* | Rock | Mbu | 1 | 5 | (5) | 0 | 9 | 0 | 14 |
| *Labeotropheus fuelleborni* | Rock | Mbu | 1 | 12 | (2) | 0 | 16 | 0 | 28 |
| (*Labeotropheus fuelleborni*) | Rock | Mbu | 2 | 0 | 0 | 0 | 7 | 0 | 7 |
| *Labeotropheus trewavasae* | Rock | Mbu | 2 | 0 | 0 | 0 | 11 | 0 | 11 |
| *Petrotilapia* sp. 'chitimba' | Rock | Mbu | 1 | (12) | 13 | 0 | 14 | 0 | 27 |
| *Labidochromis caeruleus* | Rock | Mbu | 2 | 0 | 0 | 0 | 10 | 0 | 10 |
| *Labidochromis* sp. 'hongir' | Rock | Mbu | 2 | 0 | 0 | 0 | 4 | 0 | 4 |
| *Cynotilapia zebroides* | Rock | Mbu | 2 | 0 | 0 | 0 | 21 | 0 | 21 |
| *Cynotilapia zebroides* 'Cobue' | Rock | Mbu | 1 | 13 | (11) + 1 | 0 | 18 | 0 | 32 |
| *Tropheops* sp. 'Boadzulu' | Inter | Mbu | 2 | 0 | 0 | 0 | 29 | 0 | 29 |
| *Metriaclima* (*Pseudotropheus*) *aurora* | Inter | Mbu | 2 | 0 | 0 | (14*) | 55+(14*) | (14*) | 55+(14*) |
| *Aulonocara baenschi* | Inter | B/U | 2 | 0 | 0 | (14*) | 9+(14*) | (14*) | 9+(14*) |
| *Aulonocara koningsi* | Inter | B/U | 2 | 0 | 0 | 0 | 18 | 0 | 18 |
| *Aulonocara jacobfreibergi* | Inter | B/U | 2 | 0 | 0 | 0 | 4 | 0 | 4 |
| *Copadichromis trewavasae* | Inter | B/U | 2 | 0 | 0 | 0 | 11 | 0 | 11 |
| *Copadichromis virginalis* | Sand | B/U | 1 | 18 | (11) + 1 | 0 | 15 | 0 | 34 |
| *Mchenga conophoros* | Sand | B/U | 1 | 18 | (12) | 0 | 22 | 0 | 40 |
| *Mylochromis anaphyrmus* | Sand | B/U | 1 | 14 | (4) | 0 | 0 | 0 | 14 |
| *Tramitichromis intermedius* | Sand | B/U | 1 | 18 | (10) + 1 | 0 | 19 | 0 | 38 |
| *Metriaclima* (*Pseudotropheus*) *livingstonii* | Sand | Mbu | 2 | 0 | 0 | 0 | 10 | 0 | 10 |
| *Metriaclima callainos* | Rock | Mbu | 2 | 0 | 0 | (14*) | (14*) | (14*) | (14*) |
| *Metriaclima lombardoi* | Rock | Mbu | 2 | 0 | 0 | (14*) | (14*) | (14*) | (14*) |
| *Metriaclima pyrronotos* | Rock | Mbu | 2 | 0 | 0 | (14*) | (14*) | (14*) | (14*) |
| **TOTALS** |  |  |  | 110 | (67) + 4 | 70 | 341+(70*) | (70*) | 455+(70*) |

**Supplementary Figure 4.**
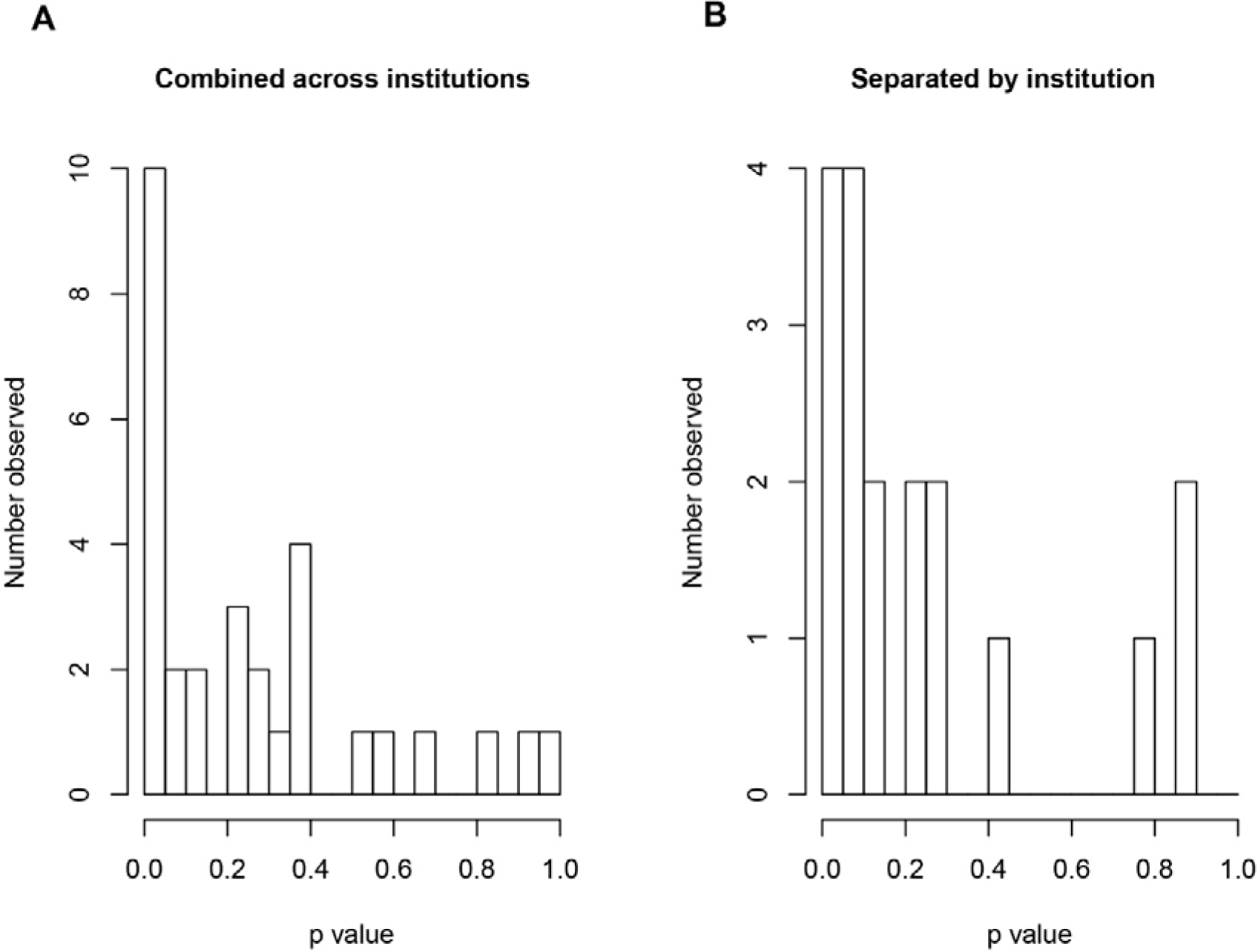
Distributions of p-values for open field microhabitat-behavior and radiation-behavior relationships in (A) primary analyses of data combined across institutions (see Supplementary Methods and Results, “Multiple hypothesis testing”), and (B) supplementary analyses of subsets of data separated by institution (see Supplementary Methods and Results, “Effects of Test Site”). For primary analyses of open field data combined across test sites, a total of 30 relationships were tested. The distribution of the resulting 30 p-values was extremely leftward skewed, illustrated by a high proportion of p-values falling below 0.05 (A; 10/30 p-values below 0.05, ∼7x more than expected by chance). Although our models controlled for test site, we also assessed whether these signals were present in the smaller “within institution” subsets of the data. Relative to the combined dataset, we had more limited statistical power to detect these relationships in the smaller subsets, so a total of 18 relationships were tested using more simplified linear mixed effects models (see “Supplementary Methods and Results” subsection “Effects of Test Site” above). Similar to the primary analyses, the distribution of the resulting p-values was leftward skewed (B). 4/18 p-values (∼22%) fell below 0.05 (two from INSTITUTION 1, two from INSTITUTION 2), and 8/18 (∼44%) fell below 0.10 (three from INSTITUTION 1, five from INSTITUTION 2), ∼4x more than expected by chance. The vast majority (7/8, ∼88%) of these relationships were matched by statistically significant (p<0.05) relationships in the primary analysis of the combined data.

